# Role of Apyrase in Mobilization of Phosphate from Extracellular Nucleotides and in Regulating Phosphate Uptake in Arabidopsis

**DOI:** 10.1101/2023.08.18.553880

**Authors:** Robert D. Slocum, Huan Wang, Alexandra A. Tomasevich, Greg Clark, Stanley J. Roux

**Affiliations:** Department of Biological Sciences, Goucher College, Towson, MD 21204; Department of Molecular Biosciences, The University of Texas at Austin, Austin, TX 78712, USA

**Author notes:** **Correspondence:** Robert D. Slocum.

**Keywords:** apyrase, eATP, auxin, extracellular ATP, nucleotide salvaging, PSR, phosphate starvation-responsive, RSA, root system architecture

## Abstract

Apyrase (nucleotide triphosphate diphosphohydrolase, NTPDase; EC 3.6.1.5) functions in a variety of plant growth and developmental processes and responses to pathogens, in part, by regulating extracellular ATP (eATP) concentrations. In this study, we investigated possible roles for apyrase in the recruitment of phosphate (Pi) from extracellular nucleotides in *Arabidopsis thaliana* seedlings that constitutively express *apyrase 1* (*APY1)*. Under Pi limitation, both WT and *APY1* seedlings had decreased Pi contents and a characteristic remodeling of root system architecture (RSA). This phosphate starvation response (PSR) was prevented by uptake of Pi released by metabolism of extracellular NTP, which supported normal seedling growth. In Pi-sufficient media, Pi contents of *APY1* seedlings were higher than in WT seedlings. Addition of NTP reduced the number of LR and decreased Pi contents of WT seedlings but markedly increased both LR and root hair (RH) formation and Pi contents of *APY1* seedlings. Genome-wide expression profiling revealed the repression of PSR genes in *APY1* seedlings, relative to WT seedlings, consistent with their elevated Pi contents. The expanded RSA of *APY1* seedlings was correlated with induction of >100 genes involved in regulation of auxin homeostasis, signaling and transport, which previous studies have shown to be increased when *APY1* is overexpressed. *APY1* regulation of [eNTP] and purinergic signaling may play a role in regulating this auxin response, resulting in enhanced uptake of Pi from the medium, including Pi released by eNTP metabolism.

## 1 Introduction

Phosphorous is an essential nutrient for plant growth and development and is acquired from the soil. Concentrations of inorganic P (Pi) in the soil are typically 1-2 µM (Schachtman et al., 1998) but high-affinity Pi transporters (PHT) in the root efficiently scavenge this nutrient and concentrate Pi in plant tissues to 10 mM or higher concentrations (Młodzińska and Zboińska, 2016; Raghothama, 1999). The vast majority of phosphate in soils is in organics such as nucleic acids, free nucleotides, sugar-P, phospholipids and polyphosphates (Huang et al., 2017; Plassard and Dell, 2010). Under phosphate limitation, plants secrete a variety of extracellular phosphatases, such as purple acid phosphatases, to hydrolyze Pi from organic molecules, for direct uptake (Tran et al., 2010).

In the rhizosphere, nucleic acids and nucleotides of various types represent a potentially significant source of P nutrition. Phosphate mobilization from extracellular nucleic acids through combined activities of nucleases, such as the P starvation-responsive (PSR) RNase RNS1 (Bariola et al., 1994), and ecto-phosphatase activities has been reported (Chen et al., 2000; Robinson et al., 2012). Free extracellular nucleotides (eNTP) could be another potential source of Pi and may be released by damaged or decaying cells in the soil (Lareen et al., 2016), in exudates from pathogen-infected roots (Yuan et al., 2018) or via vesicle-mediated secretion or efflux from root cells (Dark et al., 2011; Pietrowska-Borek et al., 2020). In rich organic soils, adenylate pools of up to 50 nmol/g soil have been reported and are tightly correlated with microbial biomass (Dyckmans et al., 2003). Estimates of free nucleotide concentrations are generally unavailable, but free ATP concentrations of ∼45 nM were estimated for a defined soil one week after inoculation of sterile medium with 10,000 cfu of a field flora (Thomas et al., 1999). Total nucleotides approaching micromolar concentrations may be expected to occur in native soils.

The extent to which salvaging of eNTP could help meet metabolic demands for cellular P is poorly understood. Although export of cellular ATP by a plasma membrane transporter has been reported (Rieder and Neuhaus, 2011), plasma membrane transporters for nucleotide uptake have not been identified in plants (Major et al., 2017). Thus, direct acquisition of Pi from nucleotides via intracellular metabolism is not possible. A role for secreted purple acid phosphatases in recruitment of Pi from dNTP (Wu et al., 2018) or ATP in soybean and bean (Zhu et al., 2020, Liang et al., 2010) has been demonstrated. Ecto-apyrase activities also function in nutrient salvaging from extracellular ATP (eATP). In potato, nucleobase uptake via apoplastic salvaging of ATP by an ecto-apyrase and coordinated activities of other ecto-phosphatases and a nucleoside hydrolase has been reported (Riewe et al., 2008). Apoplastic salvaging of NTP as a sole source of nitrogen was also shown to partially alleviate symptoms of N-limitation in Arabidopsis seedlings (Cornelius et al., 2012). Other studies support a role for ecto-apyrase activity in Pi mobilization from ATP. (Thomas et al., 1999) demonstrated that Arabidopsis seedlings ectopically expressing pea apyrase *psNTP9* had increased apyrase-like NTPase activity in the cell wall fraction and imported Pi from eATP.

Arabidopsis and soybean seedlings overexpressing *psNTP9* also exhibited an unusual expansion of root system architecture under conditions of Pi-sufficiency (Veerappa et al., 2019). Similar remodeling of root system architecture occurs in WT plants in response to Pi starvation, increasing root surface area for Pi scavenging from the soil (Péret et al., 2014; Abel, 2017). eNTP are known to modulate root growth and development (Clark et al., 2010, Tang et al., 2003, Liu et al., 2012b, Yang et al., 2015). We hypothesize that apyrase overexpression in Arabidopsis seedlings may influence Pi acquisition directly, via increased ecto-apyrase activities and release of Pi from NTP, and indirectly, by regulating eATP levels and purinergic signaling inputs into root development or other processes that influence Pi uptake. In the current study, genome-wide expression profiling was employed to elucidate possible mechanisms by which changes in *APY1* expression may influence Pi acquisition in response to altered Pi availability and NTP supplementation in Arabidopsis seedlings.

## 2 Materials and methods

### 2.1 Growth of Arabidopsis Seedlings

Seeds were surface sterilized and plated on 1/2x Murashige-Skoog (MS) medium with phosphate (0.612 mM Pi) or without Pi (Caisson Laboratories), 0.5% (w/v) MES, adjusted to pH 5.8 with KOH. Solid medium was 0.5% [w/v] Gelrite (Scott Laboratories). Some plates were supplemented with sterile-filtered 1.25 mM mixed nucleotides (1:1:1:1 ATP, GTP, CTP,UTP; 0.25 mM each for ATP, GTP, CTP and 0.5 mM UTP; Sigma Aldrich). For root growth assays, plates were placed vertically. All plates were grown at 24°C under a 16L:8D photoperiod and full-spectrum LED grow lights (150 μmoles m^−2^ s^−1^). *Arabidopsis thaliana* ecotype Wassilewskija (Ws) wild-type and an *APY1* over-expression line in the Ws background were used in the present study.

### 2.2 Root System Architecture (RSA) Analyses

ImageJ v 1.48 (http://rsbweb.nih.gov/ij/) software was used to analyze root images of vertically-grown seedlings at 10 days and 15 days of growth. Primary and lateral root lengths were measured and the ratio of total primary:secondary root lengths was calculated. Root hair (RH) number and length were measured for the distal 0.5 cm segment of primary root tips.

### 2.3 Seedling Phosphate Content Assay

Seedlings were harvested from plates on day 15, rinsed in dH_2_O, blotted and weighed. Tissues were homogenized in 1 mL of 1N H_2_SO_4_ and extracts were incubated at 42°C overnight in capped tubes. After centrifugation (16,000 x *g*, 5 min), supernatant Pi contents were quantified using a modified assay described by (Lapis-Gaza et al., 2014). Briefly, 10 µL of the hydrolysate and 90 µL dH_2_O was vortexed with 0.9 mL of freshly-prepared Pi Color Solution (0.35% (w/v) ammonium molybdate, 1.4% (w/v) ascorbic acid in 1 N H_2_SO_4_). Following a 42°C, 60 min incubation in the dark, A_820_ was measured for samples and Pi standards (1-300 nmoles KH_2_PO_4_). Pi contents were calculated from the Pi standard curve and expressed as µmol Pi g fresh weight^−1^.

### 2.4 Construction and Characterization of the *APY1* Overexpression Lines

The full-length *AtAPY1* cDNA coding region (GenBank accession no. AF093604; (Steinebrunner et al., 2000)) was cloned into the Gateway cloning vector pH7WG2 downstream of the Cauliflower mosaic virus 35S promoter. *Agrobacterium*-mediated plant transformation of *Arabidopsis thaliana* ecotype Wassilewskija (Ws-2) and hygromycin selection of transformants was as previously described (Clark et al., 2011). *APY1* expression in independent transgenic lines over-expressing *APY1* was quantified by qRT-PCR using *APY1*-specific primers (**Table S1**). Verification of the Ws-2 genetic background in WT and transgenic lines was carried out by SNP genotyping of the AT5G42320 locus, as described by (Shao et al., 2016) using primers AT5G42320-Ws2-F, AT5G42320-Col0-F and AT5G42320-Rev (**Table S1**).

### 2.5 RNA Isolation, cDNA Synthesis and qRT-PCR Analyses

Total RNA was extracted from rapid-frozen, day 5, day 11 and day 15 seedling tissues using an RNeasy Plant Mini Kit, as per the manufacturer’s protocol (Qiagen, Valencia, California). RNA samples were treated using a TURBO DNA-Free DNase I kit to remove genomic DNA contamination prior to qRT-PCR or RNA-seq analyses. For qRT-PCR assays, RNA was reverse-transcribed using a SuperScript II kit (ThermoFisher Scientific, Carlsbad, California). qRT-PCR primers (**Table S1**) were designed using NCBI Primer-BLAST (Ye et al., 2012) so that one primer of each pair spanned an exon-exon junction, preventing amplification of gDNA. Reactions (20 µL) contained 5 μL of cDNA (1 ng/µL), 0.4 µL of each primer (10 μM), 10 μL of Power SYBR Green master mix (Applied Biosystems, USA), and 4.2 μL nuclease-free water. qRT-PCR was conducted using a ViiA7 Real –Time PCR System (ThermoFisher Scientific) as follows: 95 °C for 10 min, followed by 40 cycles of 95 °C for 30 s, 58 °C for 30 s, 72 °C for 30 s in 96-well optical reaction plates (Applied Biosystems, USA). Expression of reference gene *PP2A* (AT1G69960) was used to normalize target gene expression. Relative expression was calculated using the DDCT method (Livak and Schmittgen, 2001). Dissociation curve analyses were used to check for amplification of homogenous products, whose size was verified using agarose gel electrophoresis, following secondary PCR amplification.

### 2.6 Genome-Wide Expression Analyses

Total RNA was extracted from 15-day-old seedlings that were collected 6 hours after the beginning of the light period and treated with DNase I, as described above. cDNA library preparation and Illumina sequencing (NovaSeq 6000 SR100 platform; 100 bp single-end reads) using the Tag-seq method (Lohman et al., 2016) was carried out by the Genome Sequencing and Analysis Facility (GSAF) at the University of Texas at Austin. Reads mapping and sample quality control analyses were performed as described by (Veerappa et al., 2019) for three biological replicates each of 15-day-old seedlings of the WT and *APY1* overexpression lines grown under four different experimental conditions (±P, ±NTP). An average of 13.4 million reads were mapped to the reference genome (TAIR 10.1, Araport 11 annotation; Cheng et al., 2017) for each sample, with 81.0% mapping to genes (74.0% mapped to exons). Gene differential expression analyses were carried out using the R Bioconductor module DESeq2 (Love et al., 2014). Differentially-expressed genes (DEG) were defined as having fold-change values ≥ 1.5, up or down, relative to WT expression, and *padj* ≤ 0.05. Comparison of DEG sets by Venn diagram analyses employed the web tool InteractiVenn (http://www.interactivenn.net/). Lists of DEG were analyzed for statistically over-represented gene sets in Biological Process Gene Ontology (GO) categories, using PANTHER classification system v 17.0 (http://www.pantherdb.org). Annotated gene sets for GO terms overrepresented by ≥ 2-fold (FDR p ≤ 0.05, Fisher test) were retrieved for further analyses.

### 2.7 Statistical Analyses

Statistical significance of differences between datasets was assessed by a one-way analysis of variance (ANOVA; standard weighted means analyses, independent samples), followed by Tukey’s Multiple Comparison of Means *ad hoc* test, using Vassar Stats online resources (http://vassarstats.net/anova1u.html), or using the student’s *t*-test.

## 3 Results

### 3.1 Effects of Phosphate Limitation and NTP Supplementation on Seedling Growth and Development

Growth of 15-day-old WT and *APY1* seedlings on different media is shown in **Supplemental Figure 1**. On control medium (+P, -NTP), there were no significant differences in seedling fresh weight (FW; **Figure 1**), primary root length (**Figure 2**) or number of lateral roots (LR) per seedling (**Table 1**) in the two lines although increased root hair (RH) density and length were observed for *APY1* roots (**Figures 3A, 3B, 4**).

**Figure 1.**
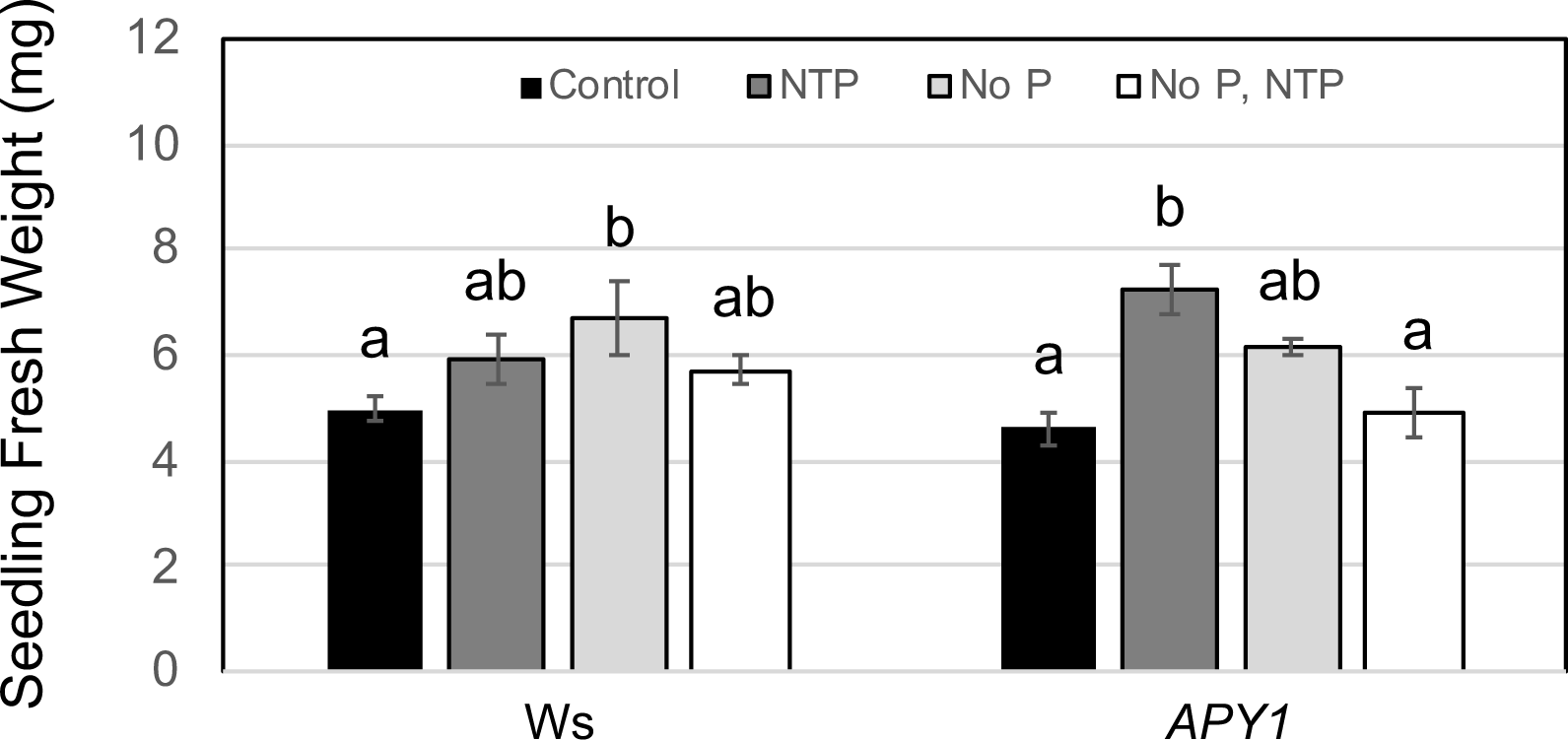
Fresh weights of 15-day-old seedlings grown in media with and without Pi, ± NTP supplementation. Data are means ± S.E. for three biological replicates of 4-6 seedlings each. Lowercase letters indicate significant difference, as determined by one-way analysis of variance (ANOVA) with *post-hoc* Tukey honest significant difference (HSD) testing (p < 0.01).

**Figure 2.**
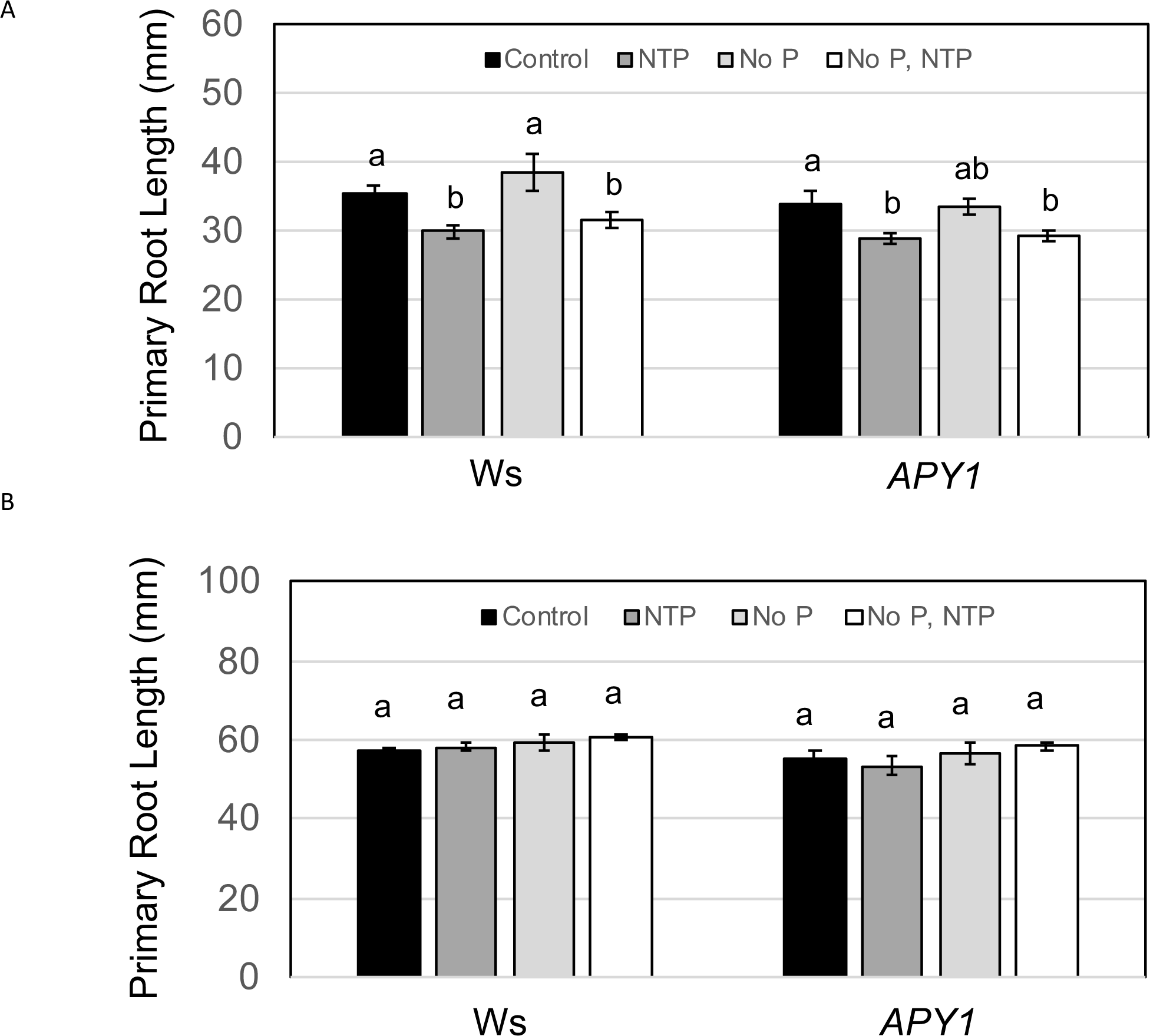
Primary root growth for day 10 seedlings (A) and day 15 seedlings (B) in response to changes in P availability and NTP supplementation. Data are means ± S.E. for 12-18 seedlings per sample (see Table 1). Lowercase letters indicate significant difference, as determined by one-way ANOVA (p < 0.01).

**Figure 3.**
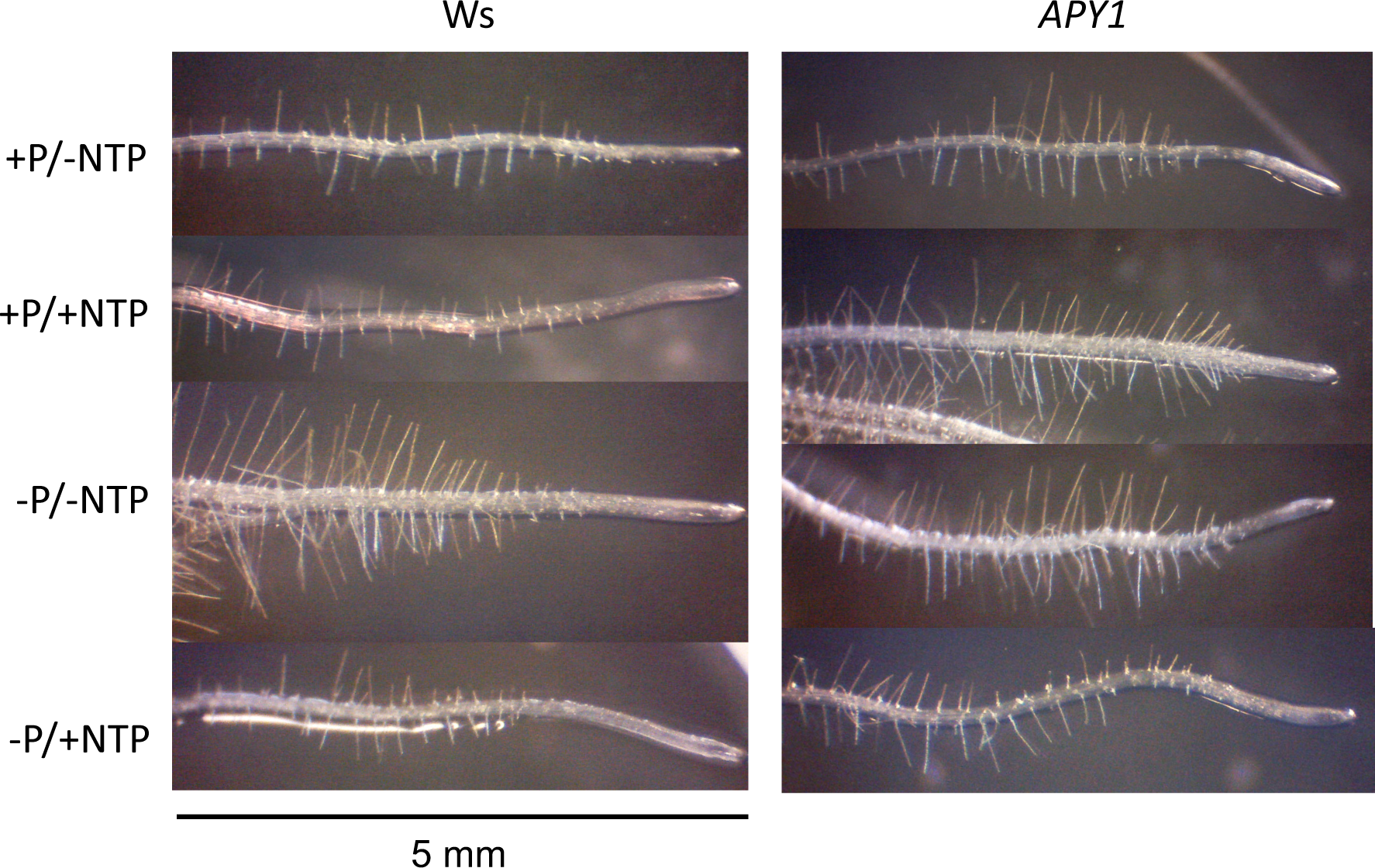
Representative photos of the distal 0.5 cm of root tips from 10-day-old Arabidopsis seedlings grown on Pi-sufficient or Pi-deficient medium with or without NTP supplementation. WT and *APY1* OE lines are shown.

**Table 1.**
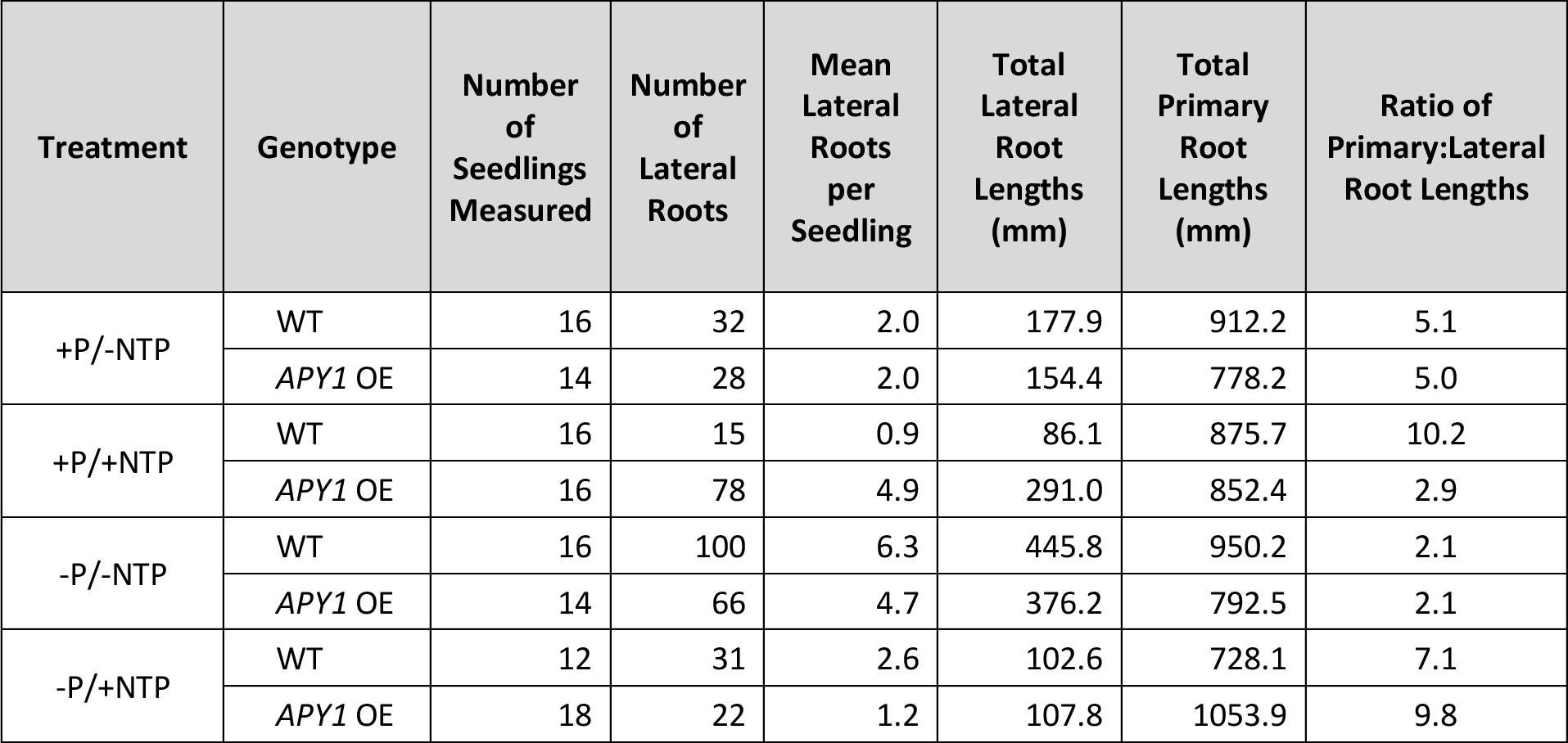
RSA remodeling in Arabidopsis seedlings in response to changes in Pi availability and NTP supplementation. Number of lateral roots per seedling and the ratio of primary:lateral root lengths in 15-day-old seedlings are indicators of RSA remodeling.

Supplementation of +P medium with NTP significantly increased the FW of *APY1* but not WT seedlings (**Figure 1**). NTP inhibited primary root growth in both lines at day 10 (**Figure 2A**) but not at day 15 (**Figure 2B**). NTP had markedly different effects on LR and RH growth in WT and *APY1* seedlings (**Figure 3**; **Table 1**). It decreased the number of LR per seedling 2-fold in WT but increased LR numbers 2.5-fold in *APY1* seedlings (**Table 1**). Total LR length was also decreased 2-fold by NTP in WT but was increased 2.5-fold in *APY1* seedlings (**Table 1**). The length of RH in WT roots was not changed by NTP supplementation, however, this treatment significantly increased RH length in *APY1* roots (**Figure 4A**). NTP supplementation decreased RH density in WT roots but increased RH density in *APY1* roots (**Figure 4B**).

**Figure 4.**
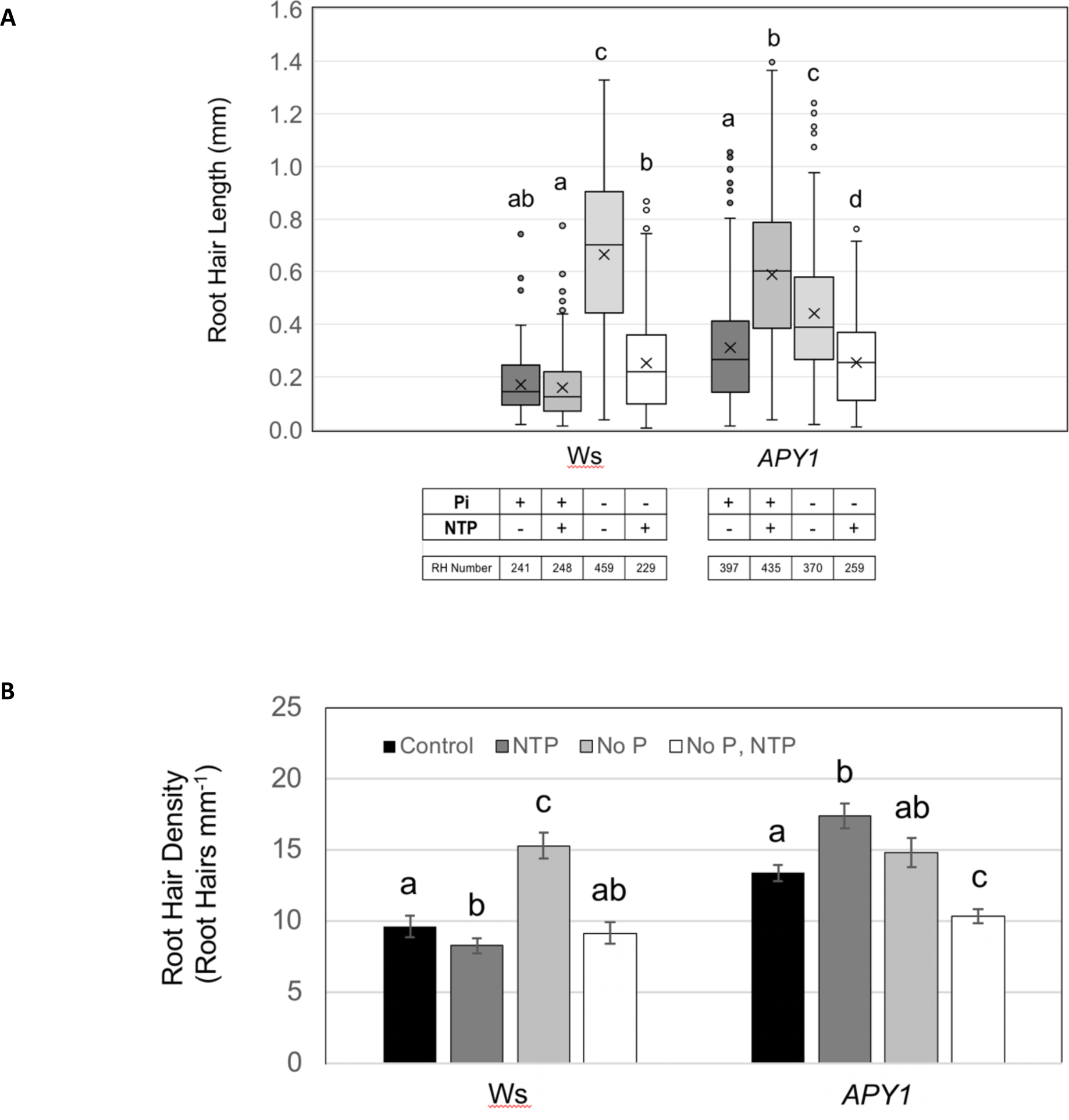
Quantification of root hair length and density in 10-day-old WT or *APY1* seedlings grown on media with or without Pi ± NTP. RH length is shown in **A**. Boxes represent 1^st^ and 3^rd^ quantiles with the center line being the median and “x” representing the mean. Whiskers are maximum and minimum values, not including outliers (>1.5 IQR). Seedling treatments and number of RH in each sample set is indicated. Root hair density is shown in **B**. Data are means ± S.E. (n ≥ 5 seedlings). Lowercase letters indicate significant difference, as determined by one-way ANOVA (p < 0.01).

The FW of WT or *APY1* seedlings growing in -P medium was not significantly different from seedlings growing in +P medium (**Figure 1**). A remodeling of RSA in response to P-limitation was observed in both WT and *APY1* seedlings growing on -P medium. The expected shortening of primary roots did not occur in either line (**Figure 2A, 2B**). Both lines had 2.5-3-fold increased numbers of LR (**Table 1**) and a 2.5-fold increase in total LR lengths (**Table 1**). RH density was not significantly different for *APY1* seedlings growing on medium with or without Pi (**Figure 4B**) but RH length was approximately half that of WT seedlings (**Figure 4A**). In contrast, WT roots exhibited a 50% increase in RH density and nearly a 4-fold increase in RH length in response to P-limitation (**Figures 3; 4A, 4B**), similar to previous observations (Ma et al., 2001).

Supplementation of -P medium with NTP did not alter seedling fresh weight (**Figure 1**). NTP decreased primary root length at day 10 (**Figure 2A**) but not at day 15 (**Figure 2B**) in both WT or *APY1* seedlings. In this medium, NTP did not affect LR growth but repressed LR initiation in both lines and the number of LR per seedling was 2-fold lower in *APY1* seedlings (**Table 1**). The RH number and lengths were markedly lower in both lines, in response to NTP, relative to seedlings growing on -P medium (**Figures 3, 4A, 4B**).

### 3.2 Pi Contents of WT and *APY1* Seedlings in Response to Changes in P Availability and NTP Supplementation

In +P medium, *APY1* seedlings showed a small but significant increase in Pi contents, compared with WT plants (**Figure 5**). NTP supplementation resulted in *APY1* Pi contents that were 2-fold higher than in WT seedlings, in part, due to a 25% decrease in Pi contents of the latter. On -P medium, the Pi contents of WT and *APY1* seedlings were not significantly different but both were reduced by 50%, relative to seedlings grown on control medium. NTP supplementation of this medium maintained WT and *APY1* seedling Pi contents that were similar to those of seedlings grown on control medium.

**Figure 5.**
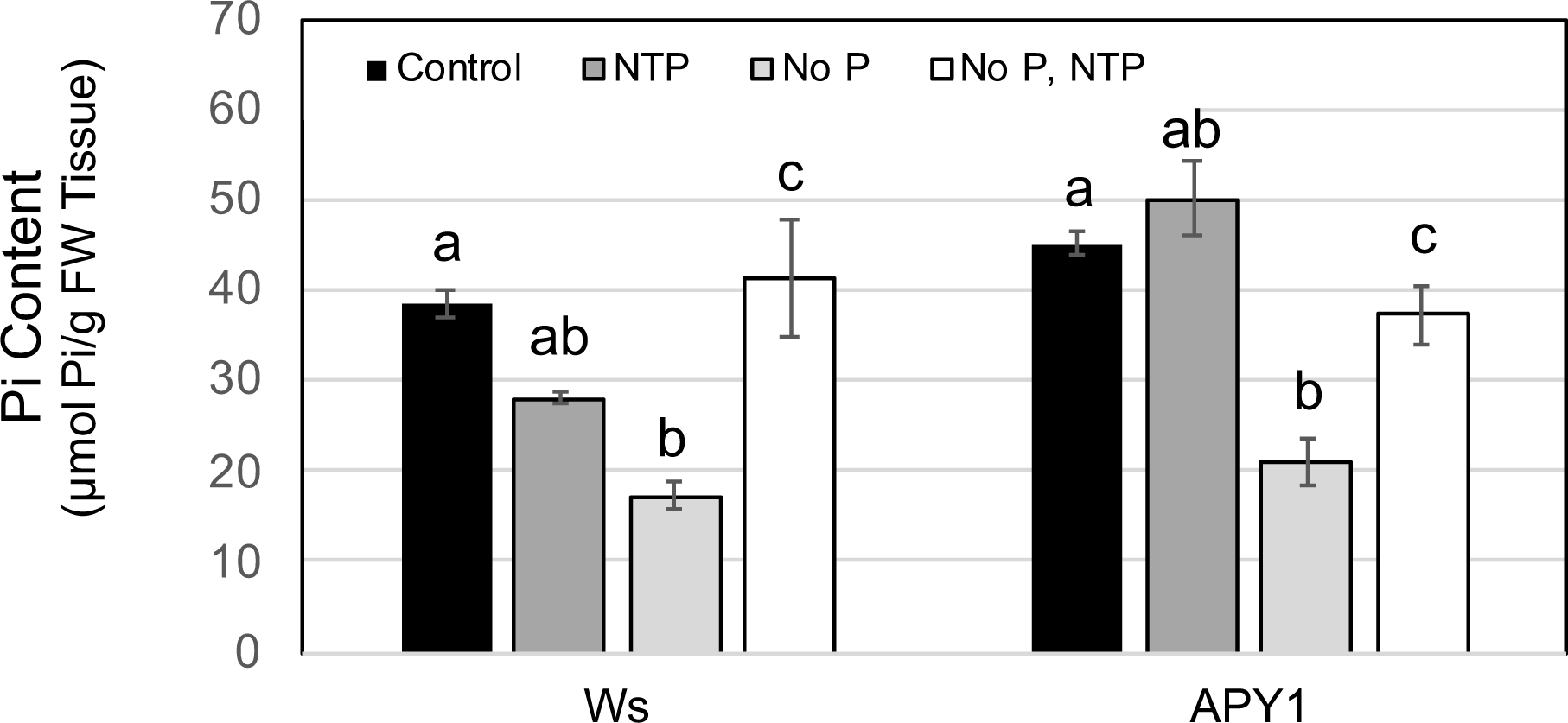
Pi contents of 15-day-old seedlings in respond to changes in Pi availability and NTP supplementation. Data are means ± S.E. of three biological replicates of 5-10 seedlings each. Lowercase letters indicate significant difference, as determined by one-way ANOVA (p < 0.01).

In preliminary studies, responses of an *apy1* line in the Ws background to changes in P availability or NTP were not different from WT seedlings (data not shown) and this line was not included in further analyses. The lack of discernible *apy1* phenotypes has been reported in previous studies (Wu et al., 2007; Liu et al., 2012b; Steinebrunner et al., 2003). *APY1* and *APY2* are functionally redundant (Wolf et al., 2007) and exhibit compensatory responses when expression of either gene is significantly reduced (unpublished data).

### 3.3 Genome-wide Expression Profiling to Investigate Responses of WT and *APY1* Seedlings to P Limitation and NTP Supplementation

Expression levels of *APY1* measured by qRT-PCR analyses were 22- to 26-fold higher than in WT seedlings at day 5, day 11 (**Supplemental Figure 2**) and day 15 stages, confirming high transgene expression in this line during seedling development. Three other independently-transformed Col-0 lines exhibited *APY1* expression levels only slightly higher than in wild-type seedlings. One transgenic line exhibited a 2.8-fold increase in *APY1* expression and a weaker Pi accumulation phenotype than the Ws line used in the current study (**Supplemental Figure 3**). Importantly, in a vertical plate assay of the Col-0 *APY1* line, 10-day-old seedlings in +P medium supplemented with NTP showed a 10-fold increase in LR number (data not shown), which is similar to the RSA phenotype observed for the Ws *APY1* line in the same medium. Genome-wide expression profiling was limited to the Ws line with the higher *APY1* overexpression.

We performed pairwise comparisons of gene expression profiles in WT and *APY1* seedlings to identify DE genes (DEG) for each of the four experimental treatments. In total, 3,780 unique genes were DE in one or more comparison groups. Fold-change expression values and annotation data were arrayed in a single matrix for comparison of expression data for individual genes, by treatment and by genotype. Sets of DEG were also subjected to Venn analyses comparing responses of the individual WT or *APY1* seedlings to different experimental treatments, or comparing WT and *APY1* seedling responses to the same treatments. In each overlap analysis, sets of genes shared by each set of DEG from a pairwise comparison, and sets of DEG unique to one or the other comparison group, were analyzed for GO BioProcess category enrichment. Genes and annotations for highly-overrepresented categories were retrieved. This approach facilitated the understanding of differences and similarities between WT and *APY1* seedlings and their responses to experimental treatments, at the transcriptome level.

Most previous studies of P metabolism have been carried out with the Arabidopsis Col-0 accessions, not the Ws ecotype used in the present study. For this reason, we first examined gene expression changes in WT and *APY1* seedlings under Pi limitation to understand how they compared with well-characterized Col-0 responses. We then investigated differences in gene expression between WT and *APY1* seedlings in the Ws background, relating to their ability to recover Pi from exogenous NTP in both P-replete and P-deficient media.

### 3.4 Responses to P-Limitation

Expression profiles for PSR genes in both WT and *APY1* seedlings were very similar to those in Col-0 seedlings (Misson et al., 2005). Among up-regulated DEG shared by WT and *APY1* seedlings, genes functioning in Pi homeostasis or cellular responses to P-starvation were enriched >65-fold and genes involved in sulfolipid or galactolipid biosynthesis were over-represented >100-fold (**Supplemental Figure 4A; Supplemental Table 2**). Catabolism of phospholipids and replacement of phospholipids with non-phosphorous galactolipids and sulfolipids is an adaptation to P-limitation (Misson et al., 2005). Sulfolipid synthesis genes *SQD1* and *SQD2* were more highly induced in *APY1* than in WT seedlings.

In both WT and *APY1* seedlings, growth on Pi-deficient media induced genes involved in sensing of cellular Pi levels and regulation of Pi homeostasis (*ITPK2, IPS1, IPS2/At4, SPX3*; **Supplemental Table 2**). Up-regulation of genes encoding secreted acid phosphatase PAP12 and inorganic Pi transporters (PHT1;1, PHT1;2, PHT1;4) would enhance extracellular scavenging and uptake of Pi (Robinson et al., 2012; Shin et al., 2004; Młodzińska and Zboińska, 2016). In *APY1* seedlings, genes encoding additional Pi transporters (PHT1;5, PHT1;9, PHO1;H1) and PHF1, a protein that increases Pi uptake by facilitating trafficking of PHT1 transporters to the plasma membrane (Gu et al., 2016), were further induced. In addition, *APY1* seedlings exhibited increased expression of genes for SPX1 and SPX2 proteins, which communicate changes in the cellular Pi status, regulating Pi homeostasis and expression of PSR genes by PHOSPHATE RESPONSE 1 (PHR1) and other core elements of the Pi regulon (Sun et al., 2016).

Accumulation of anthocyanins is a well-documented response to long-term P starvation in plants (Raghothama, 1999). In *APY1* seedlings, expression of genes encoding key enzymes regulating the synthesis of flavonones (CHS, CHI3), dihydroflavonols (F3H) and flavonols (FLS1), as well as UDP-glycosyltransferases required for anthocyanin accumulation was increased (**Supplemental Table 3**; Falcone Ferreyra et al., 2012). In contrast, *CHI* and *FLS1* were weakly induced in WT seedlings experiencing Pi deprivation. Interestingly, anthocyanidin synthesis genes (*DFR, ANS*) were not DE and increased anthocyanin levels were not detected in either WT or *APY1* seedlings, relative to those grown on +P medium.

Genes regulating root development were also enriched in both WT and *APY1* seedlings under P-limitation (**Supplemental Table 4**), consistent with previous studies that have characterized an expanded RSA as a hallmark of P-starvation (Péret et al., 2014). These included genes encoding peroxidases and other cell wall modifying enzymes (PER44, PER73, XTH14), nitrate transporter NRT2.1 and PSR and auxin-inducible genes that regulate LR and RH development.

### 3.5 Responses to NTP under P-limitation

In -P medium, relatively small numbers of genes were DE between WT and *APY1* seedlings, with or without NTP supplementation (**Supplemental Figure 5**) and both responded to P-limitation in a similar manner (section 3.3.1). However, compared with WT seedlings, genes for phospholipid remodeling (*MGD3, SQD1, SQD2, GPDP1, PLPZETA2*), Pi uptake (*PHT1;4*) and mobilization (*PPA1, PPA2, PPA4, PAP14*) and regulation of Pi homeostasis (*SPX1, SPX2*) were induced in *APY1* seedlings growing on P-deficient medium (**Supplemental Table 2**). When NTP was supplied as the sole source of P, a small number of these PSR genes remained up-regulated in *APY1* seedlings despite the fact that their Pi contents were not significantly different from those of WT seedlings and were only slightly lower than in *APY1* seedlings grown on control medium.

In -P medium, both WT and *APY1* seedlings exhibited an expanded RSA and relatively few differences in expression of genes related to root development or responses to auxin (**Supplemental Table 4**). Supplementation of this medium with NTP largely reversed this effect, consistent with Pi acquisition from NTP and the higher seedling Pi contents. A marked reduction in LR and RH development was paralleled by a near absence of DE genes regulating auxin responses in *APY1* seedlings, compared with WT seedlings.

### 3.6 Responses to NTP under conditions of P sufficiency

With and without NTP supplementation of +P media, genes related to responses to water deprivation or abscisic acid were down-regulated in *APY1*, relative to WT seedlings (**Figure 6**). Genes encoding enzymes facilitating cuticular wax synthesis and a number of plasma membrane and tonoplast aquaporins were repressed by NTP in *APY1* seedlings. Induced genes regulating indole glucosinolate biosynthesis were overrepresented in *APY1* seedlings without NTP, and this response was enhanced by NTP, likely as a result of increased *HIGH INDOLE GLUCOSINOLATE 1 (HIG1/MYB51)* expression (Gigolashvili et al., 2007). The latter response was not seen under P limitation (**Supplemental Figure 4A**), in which induction of sulfolipid synthesis genes, related to phospholipid remodeling, paralleled repression of genes for synthesis of S-containing glucosinolates.

**Figure 6.**
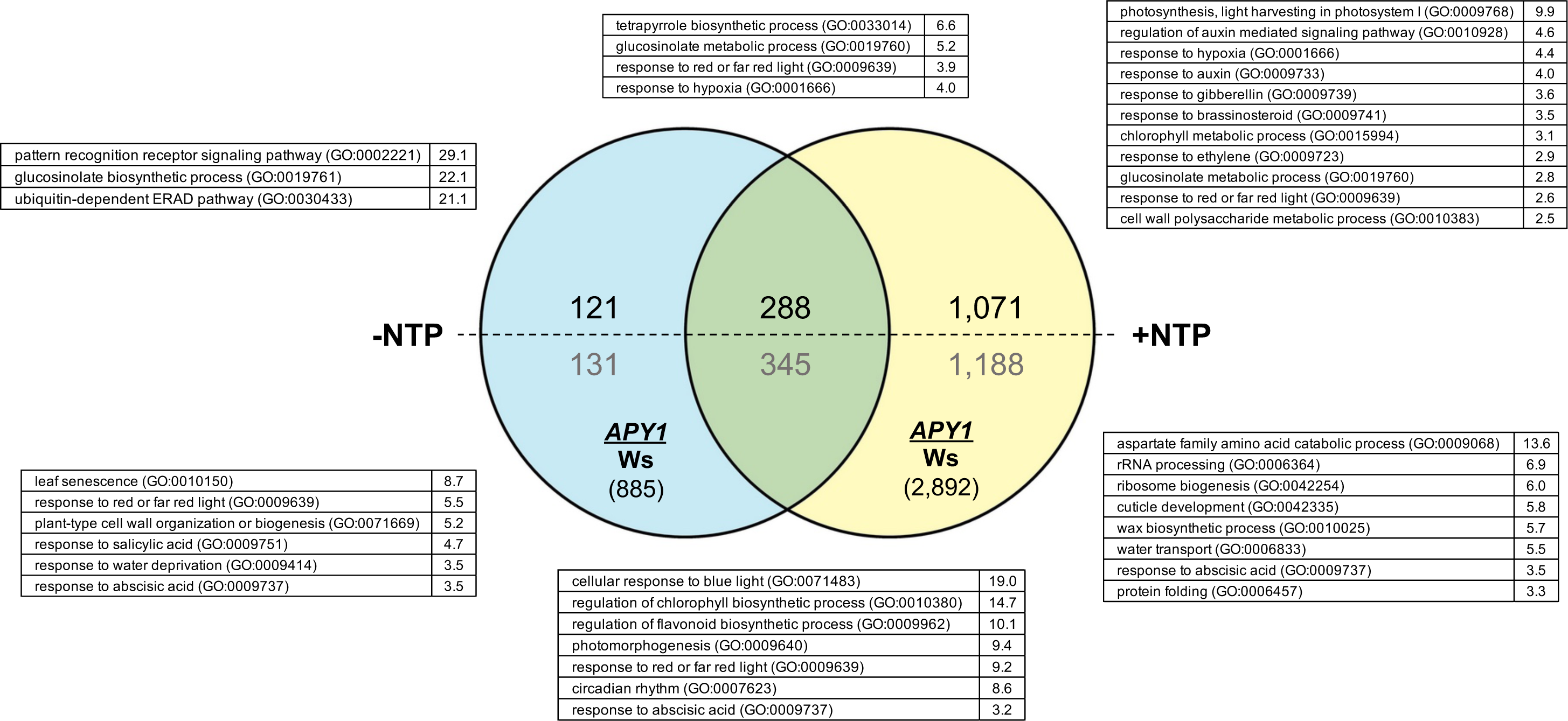
Gene enrichment analyses for 15-day-old WT and *APY1* seedlings growing under Pi-sufficient conditions, with or without NTP supplementation. Numbers of differentially-expressed genes (DEG) in each dataset are indicated in parentheses. Numbers of induced or repressed genes are indicated in black and gray, respectively. Overrepresented GO Bio Process categories and fold-enrichment values for each set of DEG are shown.

In +P medium without NTP, few genes regulating P acquisition, Pi transport or regulation of Pi homeostasis were DE between WT or *APY1* seedlings and do not offer a possible explanation for the slightly elevated Pi contents of *APY1* seedlings (**Figure 5**). However, *APY1* roots did exhibit increased RH length and density (**Figures 3B, 4**), compared with WT seedlings, but expression of genes regulating RH development was not different in *APY1* and WT seedlings (**Supplemental Table 4**). Other characteristics of RSA were not significantly different and genes regulating root development were not enriched in *APY1* seedlings (**Figure 6**). In *APY1* seedlings, genes regulating flavonoid synthesis were down-regulated, relative to WT. NTP supplementation of *APY1* seedlings in +P medium resulted in a marked repression of genes regulating flavonoid synthesis and transport (**Supplemental Table 3**).

Supplementation of the +P medium with NTP resulted in enrichment of a large number of genes involved in responses to phytohormones in *APY1* but not in WT seedlings. Induction of >100 genes regulating auxin signaling, auxin homeostasis and responses to auxin were DE (**Supplemental Table 4)**. Many of these genes regulate root development and likely contributed to the unexpected expanded RSA in these seedlings growing under P-sufficiency. These include the transcription factor UPBEAT1, which regulates the expression of a set of peroxidases that modulate the balance of reactive oxygen species (ROS) between the zones of cell proliferation and the zone of cell elongation where differentiation begins in the root (Tsukagoshi et al., 2010) and MYB73/77 and NAC001, which regulate LR development (Shin et al., 2007; Xie et al., 2000). Numerous up-regulated SAURs and cell wall remodeling enzymes (XTH22, pectin lyases, expansins) function in cell expansion growth. Remarkably, few of these changes occurred in *APY1* seedlings growing in +P medium without NTP.

The Pi contents of *APY1* seedlings grown in +P medium supplemented with NTP increased nearly 2-fold, compared with WT plants. Interestingly, apyrase 5 (*APY5*) expression in *APY1* seedlings was increased 5-fold in response to NTP in *APY1* seedlings but was not DE in WT seedlings. This was not seen in +P medium without NTP, suggesting that induction of *AtAPY5* is NTP-dependent when *AtAPY1* expression is increased. Interestingly, Xu et al. (2022) found that incuba{on of an endophy{c bacterium with Arabidopsis plants led to an increase in growth and transcriptome analyses revealed that the increase in *APY5* expression was the second highest increase in transcript abundance. Another uniquely-regulated gene in *APY1* seedlings encodes the inositol hexakisphosphate kinase VIP/VIH1 that regulates synthesis of inositol polyphosphate InsP8 as an indicator of cellular Pi status (Dong et al., 2019; Wang et al., 2021). InsP8 binding to SPX domain receptors directly or indirectly regulates Pi homeostasis (Puga et al., 2017; Ried et al., 2021). The 2-fold up-regulation of this gene by NTP is consistent with decreased transcription of PSR genes by PHOSPHATE RESPONSE 1 (PHR1), resulting from the sequestering of PHR1 in inactive complexes with SPX1/SPX2 bound to InsP8 (Puga et al., 2017). This is supported by the repression of several PHR1 target genes (**Supplemental Table 2**), including secreted acid phosphatases (*PAP8, PAP10, PAP17*), Pi transporters (*PHO1, PHT1;1, PHT1;2, VPT1*) and regulators of Pi homeostasis (*IPS1, IPS2/At4, SPX1*).

In *APY1* seedlings, NTP supplementation repressed transcription factor ZAT6, which regulates PSR genes and is induced by P-limitation (Devaiah et al., 2007). A common response to NTP in both WT and A*PY1* seedlings in +P medium was the induction of *PHO2*. PHO2 (ubiquitin-conjugating enzyme E2 24) targets PHT1 and PHO1 transporters for turnover (Liu et al., 2012a).

Gene expression data also provided insights into mechanisms of uptake and metabolism of nucleosides and nucleobases derived from NTP salvaging in the apoplast (see model, **Figure 7; Table 2**). In *APY1* seedlings, NTP supplementation of P-sufficient medium induced PM-localized purine permeases PUP14 and PUP18, which function in cellular uptake of adenine, cytosine and adenine derivatives like cytokinin (Girke et al., 2014). NTP supplementation generally repressed genes encoding enzymes of intracellular *de novo* synthesis or salvaging pathways, while nucleobase catabolism was strongly up-regulated. NTP repressed *RNS1* in both WT and A*PY1* seedlings in +P medium (**Table 2**).

**Figure 7.**
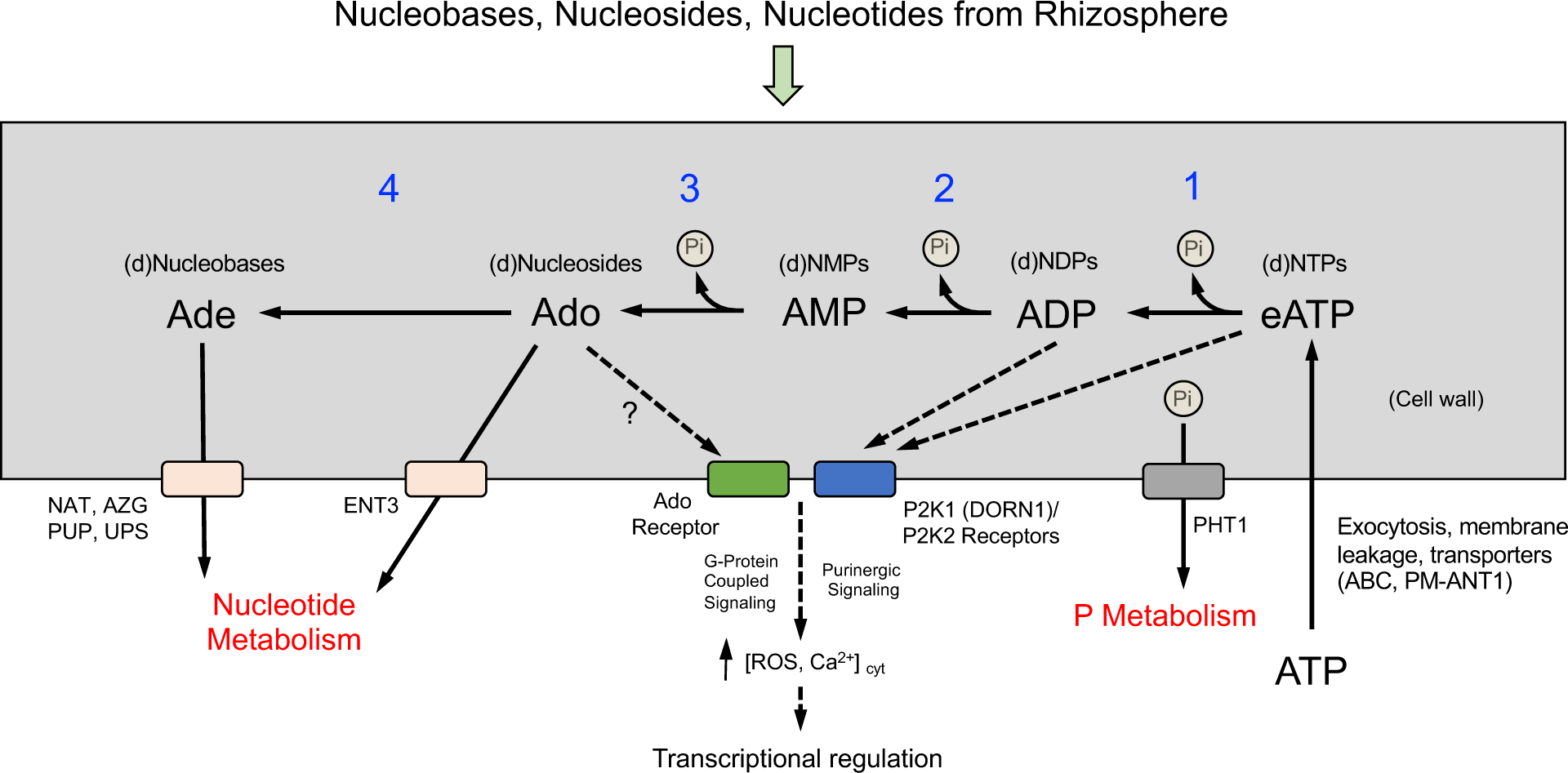
Model for apoplastic salvaging of extracellular ATP (eATP) and potential signaling pathways involving purinergic receptors or Ado receptors. Cellular and extracellular sources of eATP (and other nucleotides) are indicated. Direct uptake of nucleotides via plasma membrane NTP transporters is not known to occur. Metabolism of eATP to AMP (**steps 1, 2**) may occur via ecto-apyrase activities, with further metabolism of AMP to adenosine (Ade) by a 5’-nucleotidase activity (**step 3**), as was reported by Riewe et al. (2008). Non-specific ecto-phosphatase activities, such as purple acid phosphatases, may also metabolize eATP (**steps 1, 2, 3**; Wu et al., 2018). Pi uptake is facilitated by PM-localized PHT1 inorganic phosphate transporters. PM-localized equilibrative nucleoside transporter 3 (ENT3) imports Ado or other nucleosides into the cell (Möhlmann et al., 2010). Ado may be further metabolized to adenine (Ade) by a cell wall-localized, purine-specific nucleosidase (**step 4**; Riewe et al., 2008; Jung et al., 2011), then imported into the cell, by one or more PM-localized nucleobase transporters (Witte and Herde, 2020). Imported nucleosides and nucleobases may then enter salvaging pathways for intracellular nucleotide synthesis, or be catabolized to release nitrogen to general N metabolism (Witte and Herde, 2020). Apoplastic salvaging reactions would also regulate levels of eATP, which can bind P2K1- and P2K2-type purinergic receptors, initiating intracellular signaling cascades that regulate diverse responses in plants (Pietrowska-Borek et al., 2020). These receptors also bind ADP with high-affinity (Choi et al., 2014; Pham et al., 2020). Little is known about plant Ado receptors (Liu et al., 2021) or how Ado-mediated signaling is coordinated with eATP signaling. Initial steps of purinergic signaling (not shown) involve receptor-mediated activation of a respiratory burst NADPH oxidase RBOHD (Chen et al., 2017), leading to increased ROS in the cell wall, that can then promote cytosolic Ca^2+^ influx (Wang et al., 2019; Mohammad-Sidik et al., 2021). Plasma membrane H_2_O_2_-transporting aquaporins regulate H_2_O_2_ movement into the cell (Smirnoff and Arnaud, 2019) and may play an important role in regulating wall and cytosolic ROS levels. Increased intracellular concentrations of ROS and Ca^2+^ initiate signaling cascades resulting in transcriptional regulation of genes.

**Table 2.**
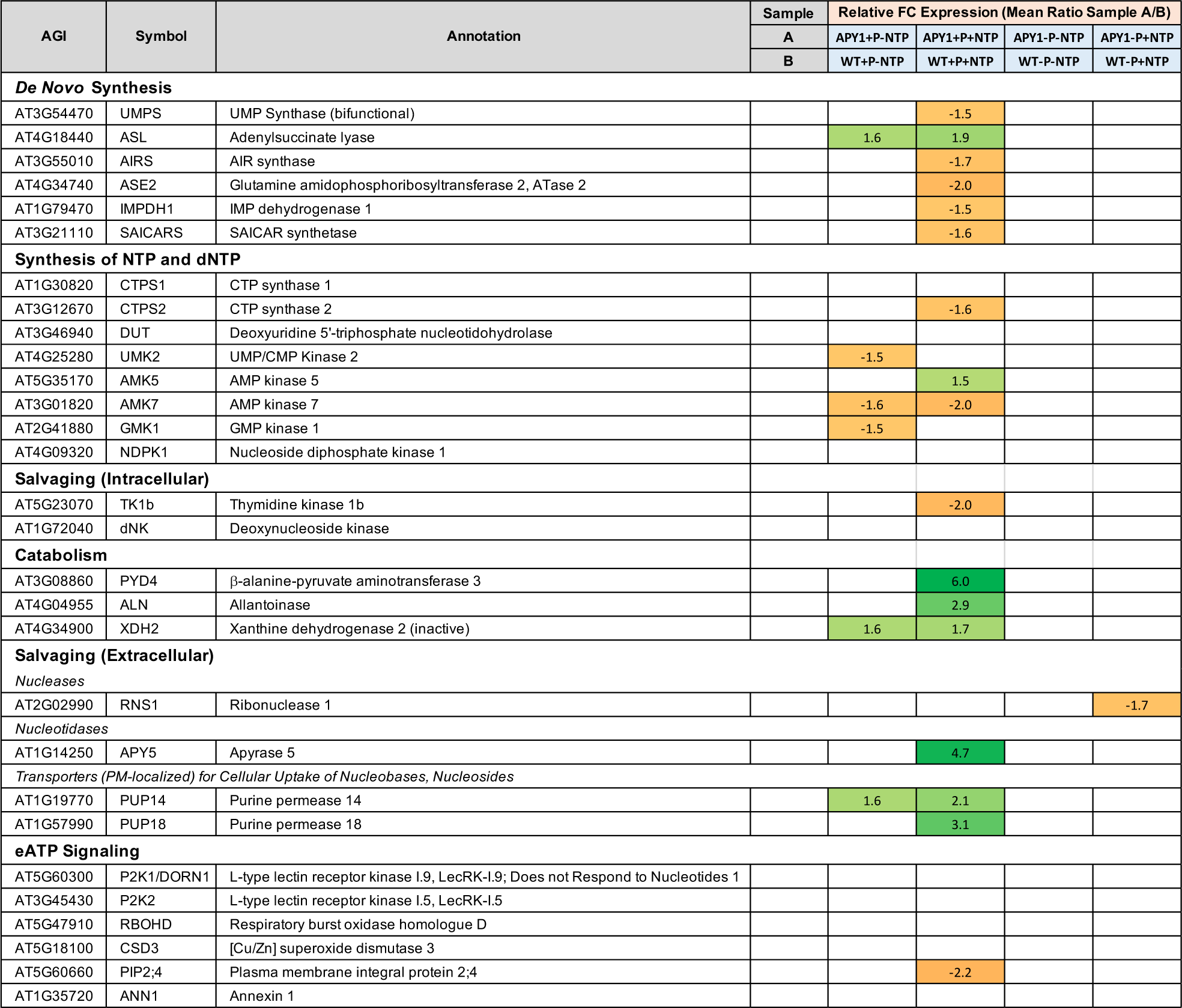
Expression of genes related to nucleotide metabolism, apoplastic salvaging of NTP, and purinergic signaling in WT or *APY1* seedlings grown on media with or without Pi, with or without NTP supplementation. The heatmap shows relative fold-change (FC) expression for each DEG.

Apart from being a source of Pi, extracellular NTP or nucleoside products of NTP salvaging could initiate signaling cascades resulting in changes in gene expression and seedling development (**Figure 7**). Differences in NTP salvaging between WT and *APY1* seedlings could alter these responses. Genes for purinergic receptors (P2K1, P2K2) and downstream signaling partners (RBOHD, ANN1) were not DE in response to NTP (**Table 2**). However, expression of H_2_O_2_-transporting PM aquaporin *PIP2;4* (Wang et al., 2020) was decreased by NTP in *APY1* seedlings but was not DE in WT seedlings.

## 4 Discussion

Arabidopsis WT and *APY1* seedlings (Ws background) generally responded to Pi-starvation as has been described for the more widely-used Col-0 accession (Misson et al., 2005). One notable difference between Col-0 versus WT and *APY1* seedlings in the present study was the lack of significant anthocyanin accumulation under Pi limitation, which has been linked to a phytochrome D mutation (*phyD-1*) in the Ws ecotype (Aukerman et al., 1997). In salt-stressed Arabidopsis seedlings, (Leschevin et al., 2021) reported significantly lower levels of anthocyanidin synthase protein in Ws seedlings, compared with Col-0 seedlings, and neither *DFR* or *ANS* were induced by Pi limitation in WT or *APY1* seedlings in the present study.

### 4.1 Metabolism of eNTP as a source of Pi in Arabidopsis seedlings

A key question addressed by this study was whether enhanced *APY1* expression could enable plants to better salvage NTPs as a source of Pi. A model for apoplastic salvaging of NTP as a source of Pi nutrition in plants, and the role for apyrase in this process, is shown in **Figure 7**. Although there are studies showing that some of the APY1 in Arabidopsis is localized in the Golgi, where it functions as an NDPase (Chiu et al., 2015; Massalski et al., 2015), there are many other types of evidence that support its additional role as an ecto-apyrase that functions in the regulation of [eATP] (Wu et al., 2007; Clark et al., 2021).

In Pi-sufficient medium, NTP supplementation had no significant effects on WT seedling growth and development or gene expression and, unexpectedly, Pi contents were slightly reduced. This may have resulted, in part, from the decreased number of LR and absorptive surface area for Pi in these plants. In contrast, the same treatment produced profound changes in *APY1* seedlings. In addition to having Pi contents that were nearly 2-fold higher than in WT seedlings, which may have resulted from increased release of Pi from NTP, the increased surface area provided by their expanded RSA would facilitate Pi uptake from the medium.

High-affinity PHT1 transporters are expressed primarily in the root epidermis and RH (Abel, 2017) and the remarkable enhancement of RH development by NTP may be the most important feature of RSA remodeling and Pi acquisition in the *APY1* seedlings. Because low [eATP] promotes RH growth and high [eATP] inhibits it (Clark et al., 2010), higher ecto-APY1 activities would help maintain lower [eATP] and thus promote RH growth. In the rice *root hairless 1 (rth1)* mutant, functional complementation with *OsAPY1* cDNA restores normal RH development (Yuo et al., 2009), further supporting an essential role of APY1 in this process.

In Pi-deficient media, one might predict that an apyrase-enhanced recovery of Pi from NTPs would result in *APY1* seedlings having higher Pi contents than WT seedlings. This was not observed. However, the gene expression profile of *APY1* seedlings grown in -P medium suggests a more robust PSR and increased potential for rapid Pi uptake and mobilization. We did not measure Pi uptake in the present study, but Thomas et al. (1999) demonstrated that identically-grown Arabidopsis seedlings ectopically expressing pea apyrase *psNTP9,* the closest pea ortholog to *AtAPY1*, exhibited a significantly higher Pi uptake than WT seedlings, during an 18 h period after transfer from low-Pi (0.1 mM Pi) to high-Pi (2 mM Pi) medium. It is possible that release and uptake of Pi from NTP occurred at a faster rate initially in *APY1* seedlings than in WT seedlings, with both acquiring approximately the same total amounts of Pi as they reached Pi homeostasis over 15 days.

Although *APY1* seedlings had elevated Pi contents in control medium (1/2x MS medium; 0.612 mM Pi), the expression of *PHT1* and *PHO1* transporter genes was not different in WT and *APY1* seedlings. The RSA of WT and *APY1* seedlings was also very similar on this medium but significantly increased RH length and density in *APY1* seedlings would provide increased surface area for Pi uptake. Again, this may have resulted from higher ecto-APY1 activities in *APY1* seedlings, maintaining lower [eATP] levels that promote RH growth (Clark et al., 2010). Interestingly, in both WT and A*PY1* seedlings in +P medium, NTP repressed the expression of *RNS1*, which is an extracellular RNase that plays an important role in mobilizing Pi from RNA under P-limitation (Bariola et al., 1994).

The salvaging of NTP from the apoplast could also impact the uptake and metabolism of nucleosides and nucleobases. Consistent with this prediction, NTP supplementation of *APY1* seedlings in +P medium induced genes for transporters and enzymes that facilitate uptake and catabolism of the nucleobase products of apoplastic NTP salvaging. While genes encoding enzymes of intracellular *de novo* synthesis or salvaging were repressed, induction of catabolic pathway activities would increase remobilization of N from purines and pyrimidines into general N metabolism (Zrenner et al., 2009; Witte and Herde, 2020). Thus, complete salvaging of nucleotides would serve to increase both P and N availability to the plant and may help explain increased *APY1* seedling fresh weights.

### 4.2 Extracellular NTP Signaling in *APY1* Seedlings Growing Under Pi-Sufficiency

Although most previous studies have examined the role of eATP on plant growth and development or stress responses, a focus of this study was Pi acquisition resulting from NTP salvaging, so a mixture of pyrimidine and purine nucleotides was provided to the seedlings to better represent the situation at a natural root/soil interface. Different responses to NTP by WT and *APY1* seedlings may be related to differences in apoplastic metabolism of eATP and purinergic signaling only, or they may further result from complex signaling mechanisms involving receptors for other extracellular nucleotides (Pietrowska-Borek et al., 2020) or their metabolites, such as Ado (Liu et al., 2021; see **Figure 7**).

Although genes for purinergic receptors (P2K1, P2K2) and downstream signaling partners (RBOHD, ANN1) were not DE in response to NTP, the plasma membrane aquaporin *PIP2;4* was repressed by NTP in *APY1* but not in WT seedlings. PIP2;4 has been shown to transport H_2_O_2_ (Wang et al., 2020) and changes in its expression might be a mechanism to regulate purinergic signaling at the level of H_2_O_2_ transport. Low (0.1 µM) concentrations of applied H_2_O_2_ stimulate and higher (>10 µM) concentrations inhibit RH growth in Arabidopsis (Clark et al., 2010). Repression of *PIP2;4* (decreased cellular uptake of H_2_O_2_), coupled with decreased H_2_O_2_ production that would result from higher ecto-APY1 activities, (decreased [eNTP] and purinergic pathway activities), may contribute to the observed increased RH growth in *APY1* seedlings.

Previous studies with Ws seedlings have reported effects of eATP on root growth and development, and several lines of evidence suggest that ecto-apyrase regulation of [eATP] and purinergic signaling impacts polar auxin transport, which is required for LR and RH formation (López-Bucio et al., 2002). Tang et al. (2003) reported that 3 mM eATP increased auxin sensitivity and disrupted basipetal auxin transport in primary root tips. They concluded that the resulting promotion of LR formation was likely due to auxin accumulation in the root tip and was regulated by eATP levels. The eATP effects on auxin transport were subsequently linked to *APY1* expression by Liu et al. (2012b), who reported an increase in basipetal auxin transport from hypocotyls into primary roots of *APY1* or *APY2* OE lines. In contrast, basipetal auxin transport in both hypocotyls and roots was decreased in the R2-4A line (*apy2* null, *APY1* suppression) but not in the single *apy1* knockout line. Chemical inhibition of apyrase activities using the specific apyrase inhibitor NGXT1913 also resulted in decreased basipetal auxin transport in gravistimulated roots (Liu et al., 2012b). These findings were consistent with the earlier results of Wu et al. (2007), who found an increase in adventitious root formation and decreased LR formation in R2-4A seedlings, possibly resulting from auxin accumulation in stem tissues.

Yang et al. (2015) further linked *APY1* expression to auxin transport. They showed that root skewing, a differential growth response that depends on local auxin gradients (Hu et al., 2021), and which can be induced either by a mechanical stimulus to the root or by locally applied ATP, was significantly increased in mutants suppressed in their expression of *APY1*. This response likely involved eATP signaling, since chemically inhibiting apyrase activity increased skewing in mutants and wild-type roots. These and other studies suggest that AtAPY1 and AtAPY2 regulate root growth and development by controlling [eATP], with altered auxin transport being one downstream response to changes in purinergic signaling. It is noteworthy that expression of these apyrases in light-grown seedlings is highest in root tip tissues where PIN auxin efflux carriers are also highly expressed (Wu et al., 2007).

Based on these studies, we predicted that NTP supplementation of WT seedlings growing in +P medium would decrease basipetal auxin transport into primary roots. This change would reduce primary root growth and auxin-regulated LR formation, which was observed. However, transcriptome data showed that few genes involved in auxin synthesis, transport, signaling or responses were DE in these seedlings, suggesting that the WT response to NTP was not primarily auxin-regulated.

Conversely, increased ecto-apyrase activities in *APY1* seedlings would be expected to reduce [eNTP], thus increasing basipetal auxin transport into the roots and promoting LR formation, which was also observed. In *APY1* seedlings, large numbers of genes involved in auxin-regulated growth were induced. Notably, the coordinated down-regulation of genes for synthesis of flavonoids, which include polar auxin transport inhibitors like quercitin (Brown et al., 2001), and increased expression of polar auxin transport components (AUX1, PIN4, PIN7) are consistent with enhanced auxin transport into the root. In the root tip, these auxin transporters play major roles in re-directing auxin flow (basipetal transport) into pericycle tissues, the site of LR initiation. They also help maintain auxin gradients that regulate root growth (Petrášek and Friml, 2009). These findings, supporting earlier work, help to explain how overexpression of *APY1* may lead to an expanded RSA for Pi acquisition, further increasing Pi contents of these seedlings under non-P-limiting conditions. A similar role for auxin in RSA remodeling has been proposed for Arabidopsis seedlings ectopically overexpressing *psNTP9* (Veerappa et al., 2019).

NTP supplementation of *APY1* seedlings growing under P-limitation did not result in changes in auxin gene expression, and LR formation was suppressed in both WT and *APY1* seedlings. Expansion of RSA under P-limitation results, in part, from increased auxin sensitivity in root tissues (López-Bucio et al., 2002), thus the slightly greater inhibition of LR formation in *APY1* plants supplemented with NTP may result from accumulation of inhibitory levels of auxin in root tissues.

We have shown that, under Pi deficiency, both WT and *APY1* plants can efficiently recruit Pi from NTP as a sole source of P, supporting normal seedling growth and development over time. In native soils, in which NTP are likely to be present at micromolar concentrations, or lower, it is unclear to what extent Pi release during NTP salvaging may contribute to overall P nutrition in plants. However, under Pi-limitation, expression profiles for PSR genes suggest that *APY1* seedlings may scavenge Pi more efficiently than WT seedlings, similar to Pi-starved *psNTP9* overexpressing Arabidopsis seedlings, upon Pi supplementation (Thomas et al., 1999). The unexpected, but potentially more important, finding of this study is that, relative to WT, the Pi contents of *APY1* seedlings were increased under conditions of Pi-sufficiency, both in the presence and absence of NTP.

Genome-wide expression profiling in the present study extends our understanding of the results of previous studies and supports a role for APY1 in modulating Pi acquisition directly, via release of Pi from NTP. These data also indicate that APY1 can impact plant Pi content through regulation of [eNTP], purinergic signaling activities, and downstream transcriptional modules that control RSA through changes in auxin transport and other processes that influence root development. These findings have potential application to the development of crops with enhanced fertilizer use efficiency and yields. Under field conditions, with periods of both limiting Pi and increased Pi availability following fertilizer applications, transgenic crops overexpressing *AtAPY1* would be expected to scavenge more Pi, potentially increasing phosphate use efficiency and limiting environmental degradation due to P leaching from soils (Veneklaas et al., 2012).

## Data Availability Statement

RNA-seq reads data have been deposited in the NCBI Sequence Read Archive (BioProject accession #PRJNA886722).

## Author Contributions

A.T. contributed to Pi content assays. H.W. provided APY1 expression and genotyping data. S.J.R. and G.C. developed the APY1 overexpression line and analyzed data. R.D.S. performed remaining experiments and analyzed transcriptome data.

## Acknowledgements

We thank the Genomic and Sequencing Facility at the University of Texas for RNA-seq support services.

