## Supplemental Figures and Tables for "Role of Apyrase in Mobilization of Phosphate from Extracellular Nucleotides and in Regulating Phosphate Uptake in Arabidopsis"

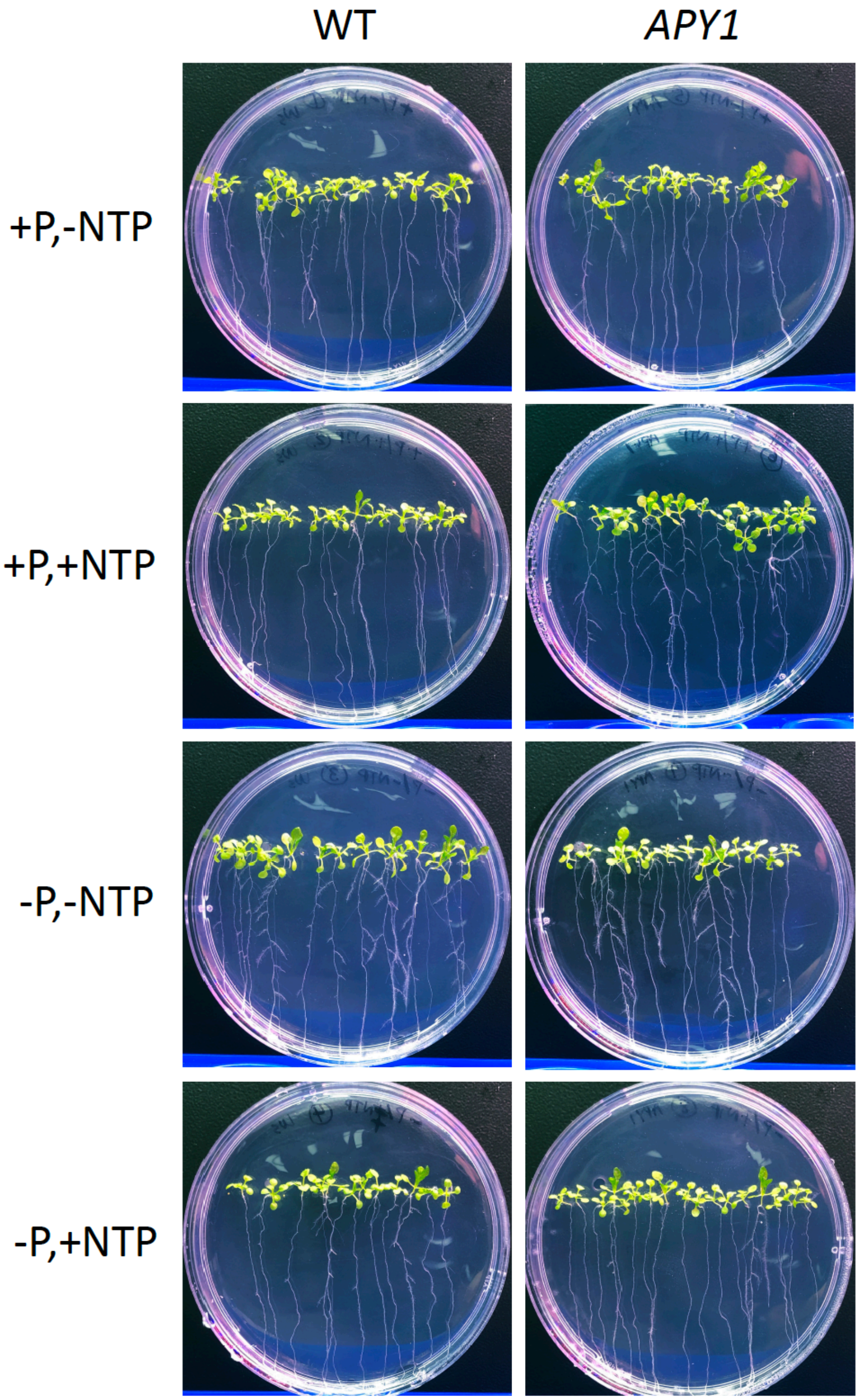


**Supplemental Figure 1.** Arabidopsis seedling RSA in response to changes in P availability, with or without NTP supplementation. Vertically-grown 15-day-old seedlings.

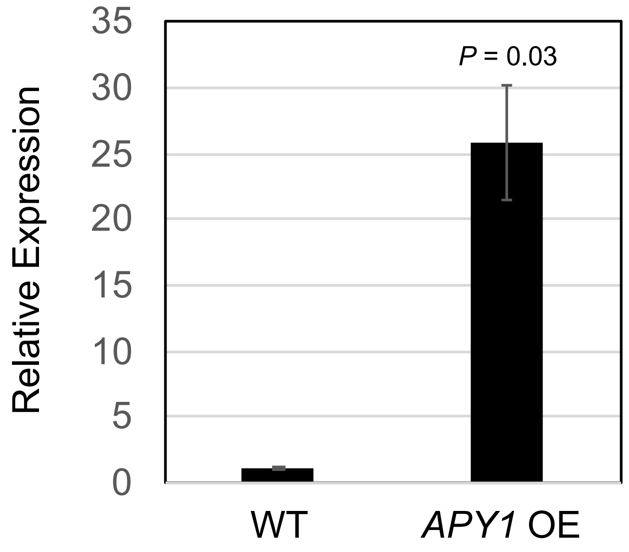


**Supplemental Figure 2.** qRT-PCR analysis of *APY1* expression in WT and *APY1* OE lines in 11-day-old Arabidopsis seedlings.  Data are means ± S.E. of 3 biological replicates of 10-15 seedlings, 3 technical replicates each. Student's *t*-test. The 318 bp *APY1* product was verified by agarose gel electrophoresis following secondary PCR amplification.

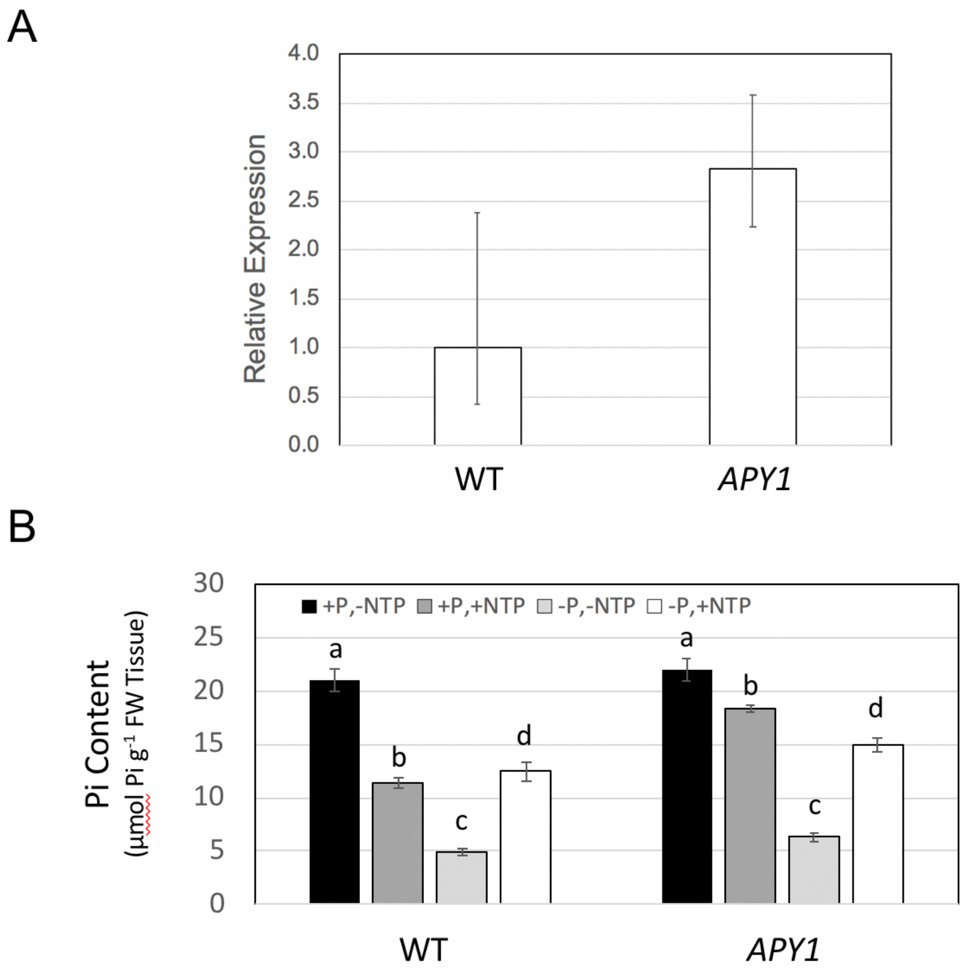


**Supplemental Figure 3.** Pi contents of an independently transformed *APY1* line (Col-0) in response to ±P, ± NTP. qRT-PCR analysis of *APY1* expression in 7-day-old WT and *APY1* lines (A). Pi contents of day 10 seedlings (B). Lowercase letters indicate significantly different groups, as determined by 1-ANOVA ($p < 0.01$). Assays were as described in the Methods section.

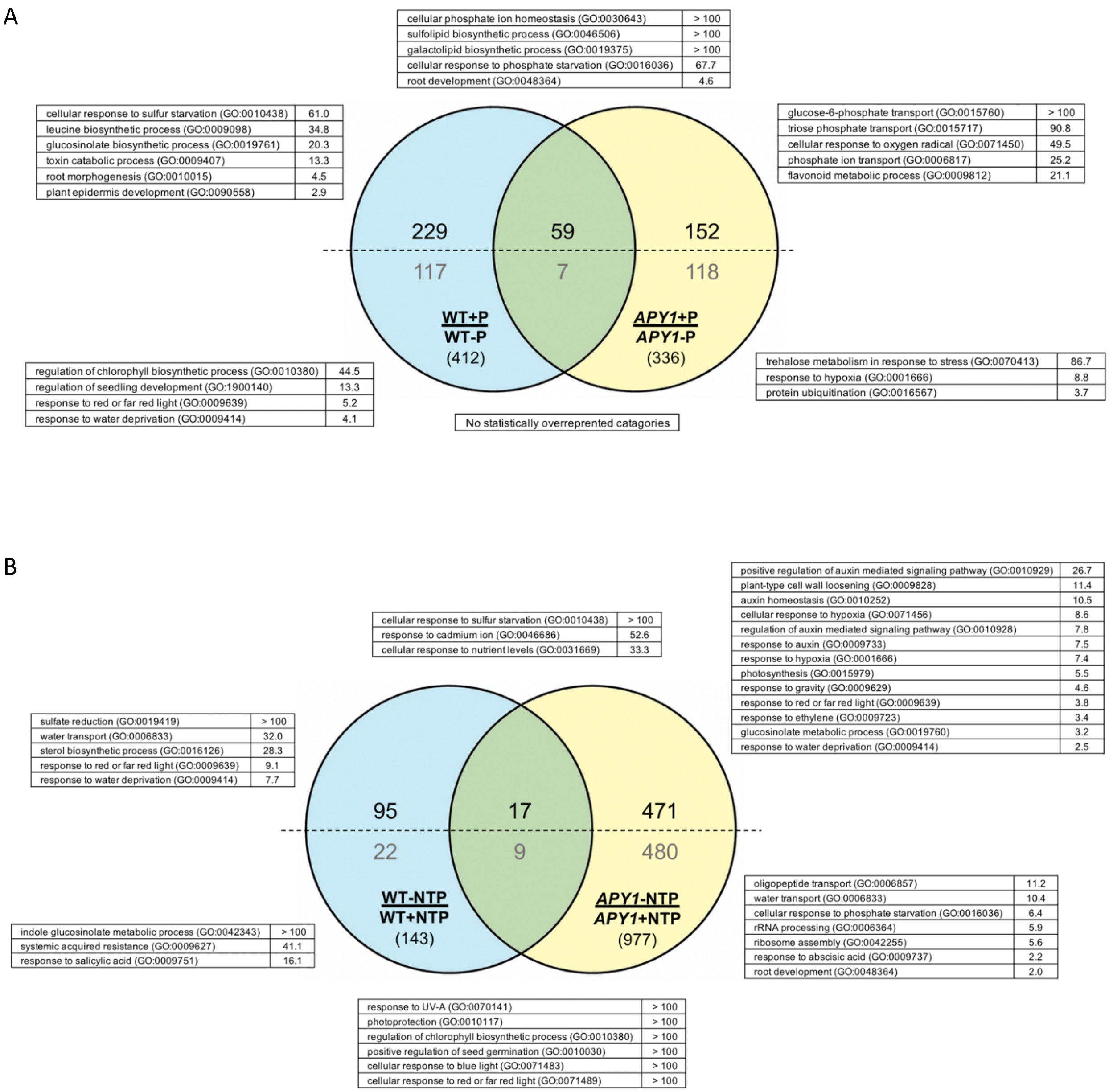


**Supplemental Figure 4.** Gene enrichment analyses for WT and *APY1* seedlings in response to Pi deficiency or NTP. 15-day-old seedlings were grown on Pi-deficient or Pi-sufficient medium, without NTP supplementation (**A**) or on Pi-replete medium, with or without NTP supplementation (**B**). Numbers of DEG in each dataset are indicated in parentheses. Numbers of induced or repressed genes are indicated in black and gray, respectively. Overrepresented GO Bio Process categories and fold-enrichment values for each set of DEG are shown.

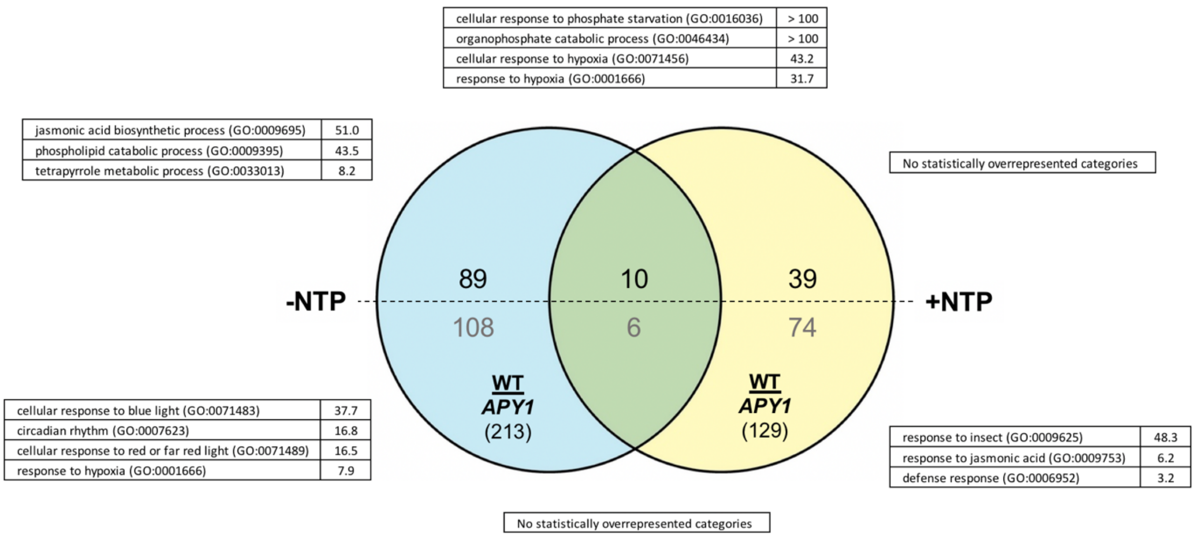


**Supplemental Figure 5.** Gene enrichment analyses for WT and *APY1* seedlings growing on Pi-deficient medium, with or without NTP supplementation. Numbers of DEG in each dataset are indicated in parentheses. Numbers of induced or repressed genes are indicated in black and gray, respectively. Overrepresented GO Bio Process categories and fold-enrichment values for each set of DEG are shown.

**Supplemental Table 1**. Primers used for verification of WT and *APY1* lines by PCR genotyping and qRT-PCR quantification of *APY1* expression.

| Primer Name | Sequence (5'->3') |
|---|---|
| Ws Genotyping | |
| AT5G42320-Col0-F | CAAATATGGTTACCATTTCTTCGTTCACAAAGAACT |
| AT5G42320-Ws2-F | AAATATGGTTACCATTTCTTCGTTCACAAAGATGA |
| AT5G42320-Rev | TTTTGACAGATGATTCCTTTTCTTCCAGACTTTT |
| qRT-PCR | |
| PP2A-F | GTTGTGTGAGCACGCAAAGA |
| PP2A-R | ACCCTCGATCAACATAATCACCC |
| APY1-F1 | CGTCCGTCTCTGTCATCGAG |
| APY1-R1 | CAGTTGCCCCAACTCTGACA |

**Supplemental Table 2.** Expression of genes related to phosphate starvation responses and regulation of Pi homeostasis in WT or *APY1* seedlings grown on media with or without Pi, ± NTP supplementation. DEG for each pairwise comparison indicate responses of either WT or *APY1* seedlings to different experimental conditions, or WT versus *APY1* responses to the same experimental treatment. The heatmap shows relative fold-change (FC) expression for each gene.

| AGI | Sample | Relative Expression (FC) | | | | | | | | Symbol | Annotation 1 |
|---|---|---|---|---|---|---|---|---|---|---|---|
| | A | WT+P-NTP | APY1+P-NTP | WT+P-NTP | APY1+P-NTP | WT+P-NTP | WT+P+NTP | WT-P-NTP | WT-P+NTP | | |
| | B | WT-P-NTP | APY1-P-NTP | WT+P+NTP | APY1+P+NTP | APY1+P-NTP | APY1+P+NTP | APY1-P-NTP | APY1-P+NTP | | |
| **Phosphate Transport** | | | | | | | | | | | |
| AT3G47420 | | | 2.1 | | -1.6 | | -2.6 | | | G3Pp1 | Glycerol-3-phosphate permease 1 |
| AT4G25220 | | 2.0 | | | | | | | | G3Pp2 | Glycerol-3-phosphate permease 2 |
| AT4G17550 | | | | | | -1.5 | -1.9 | | | G3Pp4 | Glycerol-3-phosphate permease 4 |
| AT3G52190 | | | 1.6 | | | | | | | PHF1 | Phosphate transporter traffic facilitator1, PHF1 |
| AT3G23430 | | | | | | | -1.5 | | | PHO1 | PHOSPHATE 1 |
| AT1G68740 | | | 2.1 | | | | | | | PHO1-H1 | Phosphate transporter PHO1 homolog 1 |
| AT1G35350 | | | | | | | 1.7 | | | PHO1-H8 | Phosphate transporter PHO1 homolog 8 |
| AT2G33770 | | | | 1.5 | 1.6 | | | | | PHO2 | Phosphate 2; ubiquitin-conjugating enzyme E2 24, UBC24 |
| AT5G43350 | | 1.5 | 2.2 | | -1.8 | | -1.5 | | | PHT1;1 | Inorganic phosphate transporter 1;1, PHT1;1 |
| AT5G43370 | | 2.4 | 5.2 | | -2.0 | | -2.3 | | | PHT1;2 | Inorganic phosphate transporter 1;2, PHT1;2 |
| AT2G38940 | | 1.7 | 1.8 | | -1.8 | | | 1.8 | | PHT1;4 | Inorganic phosphate transporter 1;4, PHT1;4 |
| AT2G32830 | | | 1.6 | | | | | | | PHT1;5 | Inorganic phosphate transporter 1;5, PHT1;5 |
| AT1G76430 | | | 1.7 | | | | | | | PHT1;9 | Inorganic phosphate transporter 1;9, PHT1;9 |
| AT1G63010 | | | | | | | -1.5 | | | VPT1 | Vacuolar phosphate transporter 1 |
| **Pi Mobilization** | | | | | | | | | | | |
| AT1G14250 | | | | | 1.7 | | 4.7 | | | APY5 | Apyrase 5 |
| AT3G02040 | | 1.5 | 4.3 | | -3.1 | | -1.7 | 1.7 | 2.4 | GDPD1 | Glycerophosphodiester phosphodiesterase GDPD1 |
| AT4G29690 | | | | | -1.8 | | | | | NPP3 | Ecto-nucleotide pyrophosphatase / alkaline phosphodiesterase 3 |
| AT1G13750 | | 1.6 | 1.8 | | | | | | | PAP1 | Purple acid phosphatase 1, PAP1 (inactive) |
| AT1G25230 | | | -1.9 | | | 1.6 | 2.8 | | | PAP4 | Purple acid phosphatase 4, PAP4 |
| AT2G01890 | | | | | -1.7 | | -1.6 | | | PAP8 | Purple acid phosphatase 8, PAP8 |
| AT2G16430 | | | | | | | -1.5 | | | PAP10 | Purple acid phosphatase 10, PAP10 |
| AT2G27190 | | 1.7 | 1.7 | | | | | | | PAP12 | Purple acid phosphatase 12, PAP12 |
| AT2G46880 | | 3.5 | | | | | | 1.7 | | PAP14 | Purple acid phosphatase 14, PAP14 |
| AT3G17790 | | 1.9 | 2.5 | | -1.9 | | -2.4 | | 2.0 | PAP17 | Purple acid phosphatase 17, PAP17 |
| AT3G52820 | | | 2.1 | | | | | | | PAP22 | Purple acid phosphatase 22, PAP22 |
| AT4G36350 | | | 1.5 | | | | | | | PAP25 | Purple acid phosphatase 25, PAP25 |
| AT1G73010 | | 1.6 | 2.5 | | -2.7 | | | 1.8 | 2.3 | PPA1 | Inorganic pyrophosphatase 1 |
| AT1G17710 | | 2.4 | 2.6 | | -2.9 | | | 1.7 | 2.8 | PPA2 | Inorganic pyrophosphatase 2 |
| AT2G46860 | | | 1.6 | | | | | | | PPA3 | Inorganic pyrophosphatase 3 |
| AT3G53620 | | | 1.9 | | | | | 1.5 | | PPA4 | Inorganic pyrophosphatase 4 |
| AT4G01480 | | 1.5 | 1.7 | | | | | | | PPA5 | Inorganic pyrophosphatase 5 |
| AT2G02990 | | | | -1.8 | -1.9 | | | | -1.7 | RNS1 | Ribonuclease 1 |
| AT1G14210 | | 1.6 | | | | | | | | | Ribonuclease T2, cell wall (secreted) |
| AT4G29270 | | | | | | | 1.5 | | | | HAD superfamily, subfamily IIIB acid phosphatase |
| AT2G39920 | | | | | | | 2.1 | | | | HAD superfamily, subfamily IIIB acid phosphatase |
| AT5G44020 | | | -1.5 | | | | | | | | HAD superfamily, subfamily IIIB acid phosphatase |
| **Pi Signaling, Regulation of Phosphate Starvation Responses** | | | | | | | | | | | |
| AT2G38170 | | | | | | | 2.0 | | | CAX1 | Vacuolar cation/proton exchanger 1 |
| AT1G25550 | | | | | | 1.7 | 1.9 | | | HHO3 | Transcription factor HHO3 |
| AT1G13300 | | | | | -2.2 | | -1.9 | | | HRS1 | HYPERSENSITIVITY TO LOW PI-ELICITED PRIMARY ROOT SHORTENING 1 |
| AT3G09922 | | 3.3 | 1.5 | | -1.6 | | -5.5 | | | IPS1 | INDUCED BY PI STARVATION 1 |
| AT5G03545 | | 2.8 | 2.5 | -2.0 | -3.5 | | -1.9 | | | IPS2/AT4 | INDUCED BY PI STARVATION 2 |
| AT2G43010 | | | | | | 3.6 | 3.8 | 2.0 | | PIF4 | Transcription factor phytochrome interacting factor 4 |
| AT5G04190 | | | -1.6 | | | 2.3 | 3.0 | | | PKS4 | Phytochrome kinase substrate 4 |
| AT5G20150 | | | 3.9 | | -2.0 | | -2.1 | 1.8 | 2.0 | SPX1 | SPX domain-containing protein 1 |
| AT2G26660 | | | 1.8 | | | | | | | SPX2 | SPX domain-containing protein 2 |
| AT2G45130 | | 5.0 | 2.5 | | | | | 2.4 | | SPX3 | SPX domain-containing protein 3 |
| AT5G04340 | | | | | | | -2.5 | | | ZAT6 | Zinc finger protein ZAT6 |
| **Phosphate Starvation Responsive** | | | | | | | | | | | |
| AT4G35090 | | -1.6 | | | | -2.2 | -2.3 | | | CAT2 | Catalase-2 |
| AT1G20620 | | | | | | 1.7 | 2.3 | | | CAT3 | Catalase-3 |
| AT3G08040 | | | | | -1.8 | | -3.3 | | | DTX43 | DETOXIFICATION 43 |
| AT4G08950 | | | | | | 1.8 | 2.3 | | | EXO | Phosphate-responsive 1 family protein EXORDIUM |
| AT1G35140 | | | | | | | 1.8 | | | EXL1 | Phosphate-responsive 1 family protein; EXORDIUM-like 1 |
| AT5G09470 | | | 2.4 | | | | | 1.7 | | PUMP6 | Mitochondrial uncoupling protein 6 |
| AT1G53310 | | | 1.5 | | | | | | | PPC1 | Phosphoenolpyruvate carboxylase 1 |
| AT4G02270 | | 2.5 | 1.5 | | -1.8 | | | | | RHS13 | Root hair specific 13 |
| AT3G15990 | | | | | | | -1.9 | | | SULTR3;4 | Sulfate transporter 3.4 |
| **Inositol Pyrophosphate Synthesis** | | | | | | | | | | | |
| AT4G33770 | | 1.5 | 1.7 | | | | | | | IPTK2 | Inositol 1,3,4-triphosphate 5/6 kinase 2 |
| AT3G01310 | | | | | | | 1.8 | | | VIP/VIH1 | Inositol hexakisphosphate kinase |
| **Phospholipid Remodeling** | | | | | | | | | | | |
| *Phospholipid catabolism* | | | | | | | | | | | |
| AT3G05630 | | 1.7 | 2.5 | | | | | 1.6 | | PLPZETA2 | Phospholipase D zeta 2 |
| AT3G18000 | | 2.088651826 | | | | | | | 1.5 | PEAMT1 | Phosphoethanolamine N-methyltransferase 1 |
| *Sulfolipid synthesis* | | | | | | | | | | | |
| AT5G01220 | | 2.3 | 3.1 | | | | | 1.8 | 1.6 | SQD2 | Sulfoquinovosyl transferase 2, SQD2 |
| AT4G33030 | | 1.6 | 2.1 | | | | | 1.5 | | SQD1 | UDP-sulfoquinovose synthase, SQD1 |
| *Galactolipid synthesis* | | | | | | | | | | | |
| AT5G20410 | | 2.4 | 2.6 | | | | | | | MGD2 | Monogalactosyldiacylglycerol synthase 2 |
| AT2G11810 | | 2.8 | 2.4 | | -1.6 | | | 2.4 | | MGD3 | Monogalactosyldiacylglycerol synthase 3 |

**Supplemental Table 3.** Expression of flavonoid biosynthesis genes in WT or *APY1* seedlings grown on media with or without Pi, ± NTP supplementation. DEG for each pairwise comparison indicate responses of either WT or *APY1* seedlings to different experimental conditions, or WT versus *APY1* responses to the same experimental treatment. The heatmap shows relative fold-change (FC) expression for each gene.

| AGI | Sample | Relative Expression (FC) | | | | | | | | Symbol | Annotation 1 |
|---|---|---|---|---|---|---|---|---|---|---|---|
| | A | WT+P-NTP | APY1+P-NTP | WT+P-NTP | APY1+P-NTP | WT+P-NTP | WT+P+NTP | WT-P-NTP | WT-P+NTP | | |
| | B | WT-P-NTP | APY1-P-NTP | WT+P+NTP | APY1+P+NTP | APY1+P-NTP | APY1+P+NTP | APY1-P-NTP | APY1-P+NTP | | |
| **Flavonones** | | | | | | | | | | | |
| AT5G13930 | | | 2.3 | | | -1.8 | -1.7 | | | CHS | Chalcone synthase |
| AT3G55120 | | | | | | | -1.9 | | | CHI1 | Chalcone-flavonone isomerase 1 |
| AT5G66230 | | 1.6 | | | | | | | | CHI2 | Chalcone-flavonone isomerase 2 |
| AT5G05270 | | | 2.0 | | | -1.5 | -1.5 | | | CHI3 | Chalcone-flavonone isomerase 3 |
| **Dihydroflavonols** | | | | | | | | | | | |
| AT3G51240 | | | 1.7 | | | -1.6 | -2.6 | | | F3H | Flavanone 3-hydroxylase |
| **Flavonols** | | | | | | | | | | | |
| AT5G08640 | | 1.6 | 1.7 | | | | | | | FLS1 | Flavonol synthase/flavanone 3-hydroxylase 1 |
| AT5G63600 | | | | | | | -1.5 | | -1.6 | FLS5 | Flavonol synthase/flavanone 3-hydroxylase 5 |
| **Anthocyanins** | | | | | | | | | | | |
| AT5G42800 | | Not DE | | | | | | | | DFR | Dihydroflavonol 4-reductase |
| AT4G22870 | | | | | | | | | | ANS | Anthocyanidin synthase |
| AT3G29590 | | | | | | | 2.3 | | | 5MAT | Malonyl-CoA:anthocyanidin 5-O-glucoside-6''-O-malonyltransferase |
| AT4G34135 | | | | | | | -2.4 | | | UGT73B2 | UDP-glucosyl transferase 73B2 |
| AT2G36790 | | | | | | | 1.6 | | | UGT73C6 | UDP-glycosyltransferase 73C6 |
| AT5G17050 | | | 1.5 | | | -1.9 | -2.5 | | | UGT78D2 | UDP-glycosyltransferase 78D2 |
| AT1G06000 | | | 1.6 | | | | -1.9 | | | UGT89C1 | UDP-glycosyltransferase 89C1 |
| **Regulation of flavonoid synthesis** | | | | | | | | | | | |
| AT2G26170 | | | | | | | -1.7 | | | MAX1 | MORE AXILLARY BRANCHES 1 |
| AT4G38620 | | | | | | | -1.7 | | | MYB4 | Transcription factor MYB4 |
| AT2G47460 | | | | | | | -2.0 | | | MYB12 | Transcription factor MYB12 |
| AT3G46130 | | | | | -1.8 | | -2.4 | | | MYB48 | Transcription factor MYB48 |
| AT5G49330 | | | 1.6 | | | | | | | MYB111 | Transcription factor MYB111 |
| **Flavonoid transport** | | | | | | | | | | | |
| AT4G25640 | | | | | | | -1.5 | | | DTX35 | DETOXIFICATION 35 |

**Supplemental Table 4.** Expression of genes involved in auxin metabolism, transport, signaling, responses and regulation of auxin homeostasis in WT or *APY1* seedlings grown on media with or without Pi, ± NTP supplementation. GO BioProcess categories for genes are indicated. DEG for each pairwise comparison indicate responses of either WT or *APY1* seedlings to different experimental conditions, or *APY1* versus WT responses to the same experimental treatment. The heatmap shows relative fold-change (FC) expression for each gene.

| AGI | Sample | Relative Expression (FC) | | | | | | | | Symbol | Annotation 1 | **auxin metabolic process** (GO:0009850, GO:009851, GO:0010249, GO:0010600) | **auxin homeostasis** (GO:0010252) | **auxin transport** (GO:0060918, GO:0080162, GO:0009926, GO:0010315, GO:0010328, GO:2000012) | **auxin signaling** (GO:0009734, GO:0010928) | **response to auxin** (GO:0009733) |
|---|---|---|---|---|---|---|---|---|---|---|---|---|---|---|---|---|
| | A | WT+P-NTP | APY1+P-NTP | WT+P-NTP | APY1+P-NTP | WT+P-NTP | WT+P+NTP | WT-P-NTP | WT-P+NTP | | | | | | | |
| | B | WT-P-NTP | APY1-P-NTP | WT+P+NTP | APY1+P+NTP | APY1+P-NTP | APY1+P+NTP | APY1-P-NTP | APY1-P+NTP | | | | | | | |
| AT1G28010 | | | | | | -1.7 | -3.0 | | | ABCB14 | ABC-type transporter 14 | | | | | |
| AT3G28345 | | | | | -1.8 | | -2.8 | | | ABCB15 | ABC-type transporter 15 | | | ■ | | |
| AT1G59870 | | | | -1.5 | | | | | | ABCB36 | ABC-type transporter 36 | | | ■ | ■ | ■ |
| AT2G22810 | | | | | 1.7 | | | | | ACS4 | 1-Aminocyclopropane-1-carboxylate synthase 4 | | | | | ■ |
| AT5G13300 | | | | | | | 1.5 | | | AGD3 | ARF GTPase-activating protein | | | | | ■ |
| AT4G12550 | | | | 1.7 | -2.6 | | -5.5 | | | AIR1 | Lipid-binding protein AIR1 | | | | | ■ |
| AT1G08980 | | | -1.6 | | | 1.7 | 2.0 | | | AMI1 | Amidase 1 | ■ | | | | |
| AT2G46530 | | | | | | | 1.6 | | | ARF11 | Auxin response factor 11 | | | | | ■ |
| AT3G61830 | | | | | | | 1.6 | | | ARF18 | Auxin response factor 18 | | | | | ■ |
| AT1G19220 | | | | | 1.6 | | 2.0 | | | ARF19 | Auxin response factor 19 | | | | | ■ |
| AT3G59900 | | 1.8 | | | | | | | | ARGOS | Auxin-Regulated Gene involved in Organ Size | | | | | ■ |
| AT2G38120 | | | | | | | 1.5 | | | AUX1 | AUXIN RESISTANT 1 | | | ■ | | ■ |
| AT5G12050 | | | | | 2.7 | 1.7 | 3.6 | | | BG1 | BIG GRAIN 1-like D | | | ■ | | |
| AT1G54200 | | | | | 1.7 | | 2.0 | | | BG2 | BIG GRAIN 1-like B | | | ■ | | |
| AT3G13980 | | | | | 1.5 | 1.6 | 2.2 | | | BG3 | BIG GRAIN 1-like A | | | ■ | | |
| AT5G63160 | | | | | 1.8 | -2.8 | -1.7 | -2.5 | | BT1 | BTB/POZ domain-containing protein 1 | | | | | ■ |
| AT3G48360 | | | -2.7 | -1.7 | | | 1.5 | -2.4 | | BT2 | BTB/POZ and TAZ domain-containing protein 2 | | | | | ■ |
| AT5G67480 | | 1.5 | | | | | 1.6 | | | BT4 | BTB/POZ domain-containing protein 4 | | | | | ■ |
| AT4G37610 | | | -1.7 | | | | | | | BT5 | BTB and TAZ domain protein 5 | | | | | ■ |
| AT3G55120 | | | | | | | -1.9 | | | CHI1 | Chalcone--flavonone isomerase 1 | | | | | ■ |
| AT5G13930 | | | 2.3 | | | -1.8 | -1.7 | | | CHS | Chalcone synthase; CHS | | | ■ | | ■ |
| AT3G25717 | | | | 1.6 | 1.8 | 1.8 | 2.1 | | | DVL6 | DEVIL 6 | | | | | ■ |
| AT2G30420 | | | | | | | 1.8 | | | ETC2 | ENHANCER OF TRY AND CPC 2 | | | | | ■ |
| AT3G45970 | | | -1.8 | | | | 1.9 | | | EXLA1 | Expansin-like A1 | | | | | ■ |
| AT3G55500 | | | | | | | -3.9 | | | EXPA16 | Expansin A16 | | | | | ■ |
| AT5G08640 | | 1.6 | 1.7 | | | | | | | FLS1 | Flavonol synthase/flavanone 3-hydroxylase | | | | | ■ |
| AT5G63600 | | | | | | | -1.5 | | -1.6 | FLS5 | Flavonol synthase/flavanone 3-hydroxylase 5 | | | | | ■ |
| AT5G56860 | | -1.7 | -1.5 | | | | | | | GATA21 | GATA transcription factor 21 | | | | | ■ |
| AT4G26150 | | -1.7 | | | | -2.2 | -2.0 | -1.6 | | GATA22 | GATA transcription factor 22 | | | | | ■ |
| AT4G37390 | | | | | 1.9 | | 2.6 | | | GH3.2 | Indole-3-acetic acid-amido synthetase GH3.2 | | ■ | | | ■ |
| AT2G23170 | | | | | 1.9 | | | | | GH3.3 | Indole-3-acetic acid-amido synthetase GH3.3 | | ■ | | | |
| AT4G27260 | | | | | 1.8 | | 2.1 | | | GH3.5 | Indole-3-acetic acid-amido synthetase GH3.5 | | ■ | | | ■ |
| AT5G54510 | | | | | 3.0 | | 3.8 | | | GH3.6 | Indole-3-acetic acid-amido synthetase GH3.6 | | ■ | | ■ | ■ |
| AT5G13370 | | | | | | -1.8 | -3.0 | | | GH3.12 | Indole-3-acetic acid-amido synthetase GH3.12 | ■ | | | | |
| AT5G47370 | | | | | 1.8 | | 2.2 | | | HAT2 | Homeobox-leucine zipper protein HAT2 | | | | ■ | ■ |
| AT4G16780 | | | | | 1.6 | | 2.3 | | | HAT4 | Homeobox-leucine zipper protein HAT4 | | | | | ■ |
| AT5G67060 | | | | | | | 2.2 | | | HEC1 | Transcription factor HECATE 1 | | | ■ | | |
| AT4G14560 | | | | | 2.1 | | 2.4 | | | IAA1 | AUX/IAA transcriptional regulator IAA1 | | | | | ■ |
| AT1G04240 | | | | | | | 2.1 | | | IAA3 | AUX/IAA transcriptional regulator IAA3 | | | | | ■ |
| AT1G52830 | | | | | 2.3 | | 2.4 | | | IAA6 | AUX/IAA transcriptional regulator IAA6 | | | | | ■ |
| AT3G23050 | | | | | 1.6 | | 1.6 | | | IAA7 | AUX/IAA transcriptional regulator IAA7 | | | | | ■ |
| AT1G04550 | | | | | | -1.5 | | | | IAA12 | AUX/IAA transcriptional regulator IAA12/BDL | | | | | ■ |
| AT4G14550 | | | | | 2.3 | | 1.9 | | | IAA14 | AUX/IAA transcriptional regulator IAA14 | | | | | ■ |
| AT3G15540 | | | | | 1.8 | | | | | IAA19 | AUX/IAA transcriptional regulator IAA19 | | | | | ■ |
| AT5G25890 | | | | | | | -1.8 | | | IAA28 | AUX/IAA transcriptional regulator IAA28 | | | | | ■ |
| AT4G32280 | | | | | 2.3 | | 4.8 | | | IAA29 | AUX/IAA transcriptional regulator IAA29 | | | | | ■ |
| AT1G44350 | | | | | | | 1.6 | | | ILL6 | IAA-amino acid hydrolase ILR1-like 6 | ■ | | | | |
| AT4G27280 | | | | | | | 2.3 | | | KRP1 | Calcium-binding EF-hand family protein | | | | | ■ |
| AT5G47390 | | | | | | 1.6 | 1.5 | | | KUA1 | Transcription factor KUODA1 | | ■ | | | ■ |
| AT1G08410 | | | | | | | -1.7 | | | LSG1-2 | GTPase LSG1-2 | | | ■ | | |
| AT2G26170 | | | | | | | -1.7 | | | MAX1 | MORE AXILLARY BRANCHES 1 | | | ■ | | |
| AT3G10870 | | | | | 1.5 | | 1.6 | | | MES17 | Methylesterase 17 | ■ | | | | |
| AT5G05500 | | 1.7 | | | -2.0 | | | | | MOP10 | Pollen Ole e 1 allergen and extensin family protein | | | | | ■ |
| AT2G47460 | | | | | | | -2.0 | | | MYB12 | Transcription factor MYB12 | | | | | ■ |
| AT3G28910 | | | | | | 1.6 | 2.3 | | | MYB30 | Transcription factor MYB30 | | | | | ■ |
| AT4G37260 | | | | | | | 2.5 | | | MYB73 | Transcription factor MYB73 | | | | ■ | |
| AT3G50060 | | | | | 1.5 | | 2.8 | | | MYB77 | Transcription factor MYB77 | | | | ■ | |
| AT1G56010 | | | | | -1.6 | | | | | NAC001 | NAC domain containing protein 1 | | | | ■ | |
| AT5G39610 | | | | | | | -2.0 | | | NAC006 | NAC domain containing protein 6 | | | | | ■ |
| AT5G56750 | | | | | | 1.6 | 1.8 | | 1.6 | NDL1 | N-MYC downregulated-like 1 | | | ■ | | ■ |
| AT5G11790 | | | 1.6 | | | | | | | NDL2 | N-MYC downregulated-like 2 | | | ■ | | |
| AT4G31820 | | | | | | | 1.8 | | | NPY1 | NAKED PINS IN YUC MUTANTS 1 | | | ■ | | |
| AT4G07410 | | | | | | | -1.7 | | | PCN | Transducin family protein / WD-40 repeat family protein | | | ■ | | |
| AT2G43010 | | | | | | 3.6 | 3.8 | 2.0 | | PIF4 | Transcription factor phytochrome interacting factor 4 | ■ | | | ■ | |
| AT3G59060 | | | | | | | 1.6 | | | PIF5 | Transcription factor phytochrome interacting factor 5 | ■ | | | ■ | |
| AT2G17500 | | | | | -1.8 | | -1.8 | | -1.5 | PILS5 | Auxin efflux carrier family protein; PIN-LIKES 5 | | ■ | ■ | | ■ |
| AT2G01420 | | | | | 1.5 | | 2.3 | | | PIN4 | Auxin efflux carrier component 4 | | ■ | ■ | | |
| AT1G23080 | | | | | 1.9 | | 2.7 | | | PIN7 | Auxin efflux carrier component 7 | | ■ | ■ | | |
| AT3G05630 | | 1.7 | 2.5 | | | | | 1.6 | | PLPZETA2 | Phospholipase D zeta 2 | | | | | ■ |
| AT4G39403 | | | | | | | 2.7 | | | PLS | POLARIS | | ■ | ■ | | ■ |
| AT5G48900 | | 1.5 | | | 2.0 | | 1.9 | | | PLY20 | Pectate lyase 20 | | | | | ■ |
| AT3G07010 | | | | | 1.5 | | 2.1 | | | PLY8 | Pectate lyase 8 | | | | | ■ |
| AT3G30350 | | | | | 1.7 | | 2.2 | | | RGF4 | Root meristem growth factor 4 | | | ■ | | |
| AT4G16515 | | | | | 2.8 | | 2.9 | | | RGF6 | Root meristem growth factor 6 | | | ■ | | ■ |
| AT5G64770 | | | | | 1.5 | | | | | RGF9 | Root meristem growth factor 9 | | | ■ | | ■ |
| AT5G17300 | | | -1.9 | | -1.9 | | -2.8 | -1.8 | | RVE1 | REVEILLE 1 | ■ | | | ■ | |
| AT4G34770 | | | -1.7 | | | 2.1 | 4.6 | | | SAUR1 | Auxin-responsive protein SAUR1 | | | | | ■ |
| AT4G34790 | | | | | | 1.9 | 4.1 | | | SAUR3 | Auxin-responsive protein SAUR3 | | | | | ■ |
| AT4G34810 | | 1.7 | | | | | | | | SAUR5 | Auxin-responsive protein SAUR5 | | | | | ■ |
| AT2G21200 | | | | | 2.2 | | 2.2 | | | SAUR7 | Auxin-responsive protein SAUR7 | | | | | ■ |
| AT2G18010 | | | | | | | 2.5 | | | SAUR10 | Auxin-responsive protein SAUR10 | | | | | ■ |
| AT4G38825 | | | | | 1.8 | | 5.0 | | | SAUR13 | Auxin-responsive protein SAUR13 | | | | | ■ |
| AT4G38850 | | | | | 1.8 | | 1.7 | | | SAUR15 | Auxin-responsive protein SAUR15 | | | | | ■ |
| AT4G38860 | | | | | 1.7 | | 2.0 | | | SAUR16 | Auxin-responsive protein SAUR16 | | | | | ■ |
| AT5G18010 | | | | | 1.9 | | 3.0 | | | SAUR19 | Auxin-responsive protein SAUR19 | | | ■ | | |
| AT5G18020 | | | | | | -1.8 | | | | SAUR20 | Auxin-responsive protein SAUR20 | | | | | ■ |
| AT5G18030 | | | | | 1.9 | | 2.5 | | | SAUR21 | Auxin-responsive protein SAUR21 | | | | | ■ |
| AT5G18050 | | | | | 3.0 | | 4.6 | | | SAUR22 | Auxin-responsive protein SAUR22 | | | | | ■ |
| AT5G18060 | | | | | 2.4 | | 3.3 | | | SAUR23 | Auxin-responsive protein SAUR23 | | | | | ■ |
| AT5G18080 | | | | | 2.5 | | 2.5 | | | SAUR24 | Auxin-responsive protein SAUR24 | | | | | ■ |
| AT3G03850 | | | | | 2.4 | | 2.2 | | | SAUR26 | Auxin-responsive protein SAUR26 | | | | | ■ |
| AT3G03840 | | | | | 1.9 | | 1.8 | | | SAUR27 | Auxin-responsive protein SAUR27 | | | | | ■ |
| AT3G03830 | | | | | | | 3.4 | | | SAUR28 | Auxin-responsive protein SAUR28 | | | ■ | | ■ |
| AT3G03820 | | | | | 1.8 | | 2.4 | | | SAUR29 | Auxin-responsive protein SAUR29 | | | | | ■ |
| AT5G53590 | | | | | | | -2.1 | | | SAUR30 | Auxin-responsive protein SAUR30 | | | | | ■ |
| AT4G31320 | | 1.6 | | | | | | | | SAUR37 | Auxin-responsive protein SAUR37 | | | | | ■ |
| AT2G37030 | | | | | | | 2.4 | | | SAUR46 | Auxin-responsive protein SAUR46 | | | | | ■ |
| AT4G34750 | | | | | | | 2.1 | | | SAUR49 | Auxin-responsive protein SAUR49 | | | | | ■ |
| AT4G34760 | | | | | 1.5 | | 1.7 | | | SAUR50 | Auxin-responsive protein SAUR50 | | | | | ■ |
| AT3G53250 | | | | | 2.2 | | 4.3 | | | SAUR57 | Auxin-responsive protein SAUR57 | | | | | ■ |
| AT3G60690 | | | | | | | -1.6 | | | SAUR59 | Auxin-responsive protein SAUR59 | | | | | ■ |
| AT1G20470 | | -1.6 | | | | -2.8 | -3.0 | -1.5 | | SAUR60 | Auxin-responsive protein SAUR60 | | | | | ■ |
| AT1G29440 | | | | | 1.9 | | 1.7 | | | SAUR63 | Auxin-responsive protein SAUR63 | | | ■ | | |
| AT1G29450 | | | | | 2.0 | | 2.7 | | | SAUR64 | Auxin-responsive protein SAUR64 | | | | | ■ |
| AT1G29460 | | | | | 1.8 | 1.7 | 2.7 | | | SAUR65 | Auxin-responsive protein SAUR65 | | | | | ■ |
| AT1G29500 | | | | 1.5 | 1.8 | | 1.7 | | | SAUR66 | Auxin-responsive protein SAUR66 | | | | | ■ |
| AT1G29510 | | | | | 1.7 | | 1.7 | | | SAUR67 | Auxin-responsive protein SAUR67 | | | | | ■ |
| AT5G10990 | | | | | | | -2.1 | | | SAUR69 | Auxin-responsive protein SAUR69 | | | | | ■ |
| AT5G20820 | | | | | | | 2.8 | | | SAUR76 | Auxin-responsive protein SAUR76 | | | | | ■ |
| AT1G70560 | | | | | 1.5 | | 1.8 | | | TAA1 | Tryptophan aminotransferase of Arabidopsis 1 | ■ | | | ■ | |
| AT1G05680 | | | | | | -2.0 | -3.5 | | | UGT74E2 | UDP-glycosyltransferase 74E2 | ■ | | | | |
| AT4G15550 | | | 1.5 | | | -1.8 | -3.4 | | | UGT75D1 | UDP-glycosyltransferase 75D1 | ■ | | | | |
| AT3G29370 | | | | | 2.2 | 1.7 | 3.8 | | | UPBEAT | Transcription factor UPBEAT-like protein | | | | | ■ |
| AT1G75500 | | | | | | | 1.6 | | | WAT1 | Walls Are Thin 1, WAT1 | | | ■ | | |
| AT5G57560 | | | | | | | 3.7 | | | XTH22 | Xyloglucan endotransglucosylase/hydrolase 22 | | | | | ■ |
| AT4G28720 | | | | | 1.9 | | 2.6 | | | YUC8 | Flavin-binding monooxygenase family protein | ■ | | | | |
| AT1G07135 | | | | | | | 3.1 | | | | Glycine-rich protein | | | | | ■ |
| AT4G23820 | | | | | 1.6 | | 1.9 | | | | Pectin lyase-like superfamily protein | | | | | ■ |
| AT4G14819 | | | | | 2.2 | | 2.8 | | | | Protein of unknown function (DUF1677) | | | | | ■ |
